# Parental genome unification is highly erroneous in mammalian embryos

**DOI:** 10.1101/2020.08.27.269779

**Authors:** Tommaso Cavazza, Antonio Z Politi, Patrick Aldag, Clara Baker, Kay Elder, Martyn Blayney, Andrea Lucas-Hahn, Heiner Niemann, Melina Schuh

## Abstract

The vast majority of human embryos are aneuploid. Aneuploidy frequently arises during the early mitotic divisions of the embryo, but the origin of this remains elusive. Using bovine embryos as a model for human embryos, we identify an error-prone mechanism of parental genome unification which often results in aneuploidy. Surprisingly, genome unification initiates hours before breakdown of the two pronuclei that encapsulate the parental genomes. While still within intact pronuclei, the parental genomes polarize towards each other, in a process driven by centrosomes, dynein, and microtubules. The maternal and paternal chromosomes eventually cluster at the pronuclear interface, in direct proximity to each other. Parental genome clustering often fails however, leading to massive chromosome segregation errors, incompatible with healthy embryo development. Nucleoli, which associate with chromatin, also cluster at the pronuclear interface in human zygotes. Defects in nucleolar clustering correlate with failure in human embryo development, suggesting a conserved mechanism.

## Introduction

Around 50-70% of human cleavage embryos are aneuploid – they carry an incorrect number of chromosomes (McCoy et al., 2015b; van Echten-Arends et al., 2011; Vanneste et al., 2009; Vera-Rodriguez et al., 2015). Most aneuploid embryos do not develop to term, making aneuploidy in embryos a leading cause of miscarriages and infertility (Benkhalifa et al., 2005; Fritz et al., 2001). Consistent with high aneuploidy rates in human embryos, only half of the embryos generated *in vitro* develop to the blastocyst stage (Ottolini et al., 2017), and only 30% of natural conceptions result in a live birth (Macklon et al., 2002). It is estimated that 10-20% of embryonic aneuploidy arises from chromosome segregation errors during the meiotic divisions of the egg (Lee and Kiessling, 2017; McCoy, 2017; McCoy et al., 2015b). The majority however is thought to arise during the mitotic divisions of the embryo (Lee and Kiessling, 2017; McCoy et al., 2015b). Mitotic errors have been linked to abnormal division events during early embryo development (Kort et al., 2016; Lee and Kiessling, 2017; McCoy, 2017). However, chromosome segregation has not yet been followed in live human embryos, and the cellular origin of mitotic aneuploidy remains unclear.

Chromosome segregation has been studied in early mouse embryos (Courtois et al., 2012; Mashiko et al., 2020; Reichmann et al., 2018). However, mouse embryos differ from human and other mammalian embryos in several crucial aspects: they have lower aneuploidy rates (Lee and Kiessling, 2017; Lightfoot et al., 2006), and the timing of their embryonic divisions (Niakan et al., 2012) and mechanism of developmental reprogramming are different (Fogarty et al., 2017). Moreover, early mouse embryos lack centrosomes (Courtois et al., 2012; Fishman et al., 2018; Woolley and Fawcett, 1973), which have essential functions during embryonic development in most other mammalian species. For instance, centrosomes drive the inward movement of the two pronuclei that encapsulate the parental genomes in the fertilized egg (zygote) (Malone et al., 2003; Payne et al., 2003), and are thought to promote the assembly of the first mitotic spindle (Fishman et al., 2018; Hewitson et al., 2002), the machinery that segregates the chromosomes. The importance of centrosomes for early embryo development is further supported by the identification of genetic variants of the centrosomal kinase PLK4 that have been linked to aneuploidy in human embryos (McCoy et al., 2015a).

Studies in human zygotes are limited by ethical considerations, the lack of available biological material, and legal restrictions. Bovine embryos closely resemble human embryos in their development: they contain centrosomes (Fishman et al., 2018; Navara et al., 1994), display similar timings of early embryonic divisions (Lequarre et al., 2003) and rates of aneuploidy (Destouni et al., 2016; Lee and Kiessling, 2017), and are reprogrammed by related mechanisms (Simmet et al., 2018).

In this study, we established a high-resolution live cell imaging system for bovine embryos to identify potential causes of errors during early mammalian embryogenesis. Using this system, we found that chromosome segregation errors frequently arise from defects during parental genome unification. We report that the unification of the maternal and paternal genomes initiates while each genome is still in a separate nucleus, much earlier than had been previously assumed. The parental genomes cluster in close proximity to each other at the pronuclear interface, which depends on centrosomes being located at the pronuclear interface. However, clustering often fails leading to chromosome segregation errors which preclude the development of a healthy embryo. This crucial role of chromosome clustering is likely to be conserved in human zygotes, as we found that nucleolar clustering correlates with normal human blastocyst development. Together, our work suggests defects during parental genome clustering as a major contributor to the high rates of aneuploidy in human and bovine embryos.

## Results

### Parental genome unification initiates in intact nuclei upon fertilization

We established a high-resolution live cell imaging system for bovine embryos as a model system for human embryos (see STAR Methods; Movie S1). We confirmed that imaged embryos developed into blastocysts with similar efficiencies as non-imaged embryos (Figure S1A). Thus, the imaging system did not interfere with embryo development.

We then used our imaging system to study the first mitotic division in the zygote (Movie S1). Strikingly, we frequently observed chromosome segregation errors upon unification of the maternal and paternal chromosomes on the first mitotic spindle (Movie S1), as we outline in detail below. This prompted us to investigate how these errors arise as the parental genomes are united. Upon fertilization, the parental genomes were first packaged into two separate pronuclei in the periphery of the zygote. The pronuclei subsequently migrated inwards and the sperm centrosome was duplicated. During this migration, the chromatin adopted an unusual configuration: instead of being homogeneously distributed throughout the pronuclear volume, it became clustered and polarized inside intact pronuclei (Figures 1A and 1B). The chromatin was highly concentrated in the direction of migration, so that when the pronuclei congressed, chromatin was clustered at the pronuclear interface (Figures 1B-1D and S1B). The chromosomes in both pronuclei were already in close proximity to each other when the male and female pronuclei reached juxtaposition.

**Figure 1.**
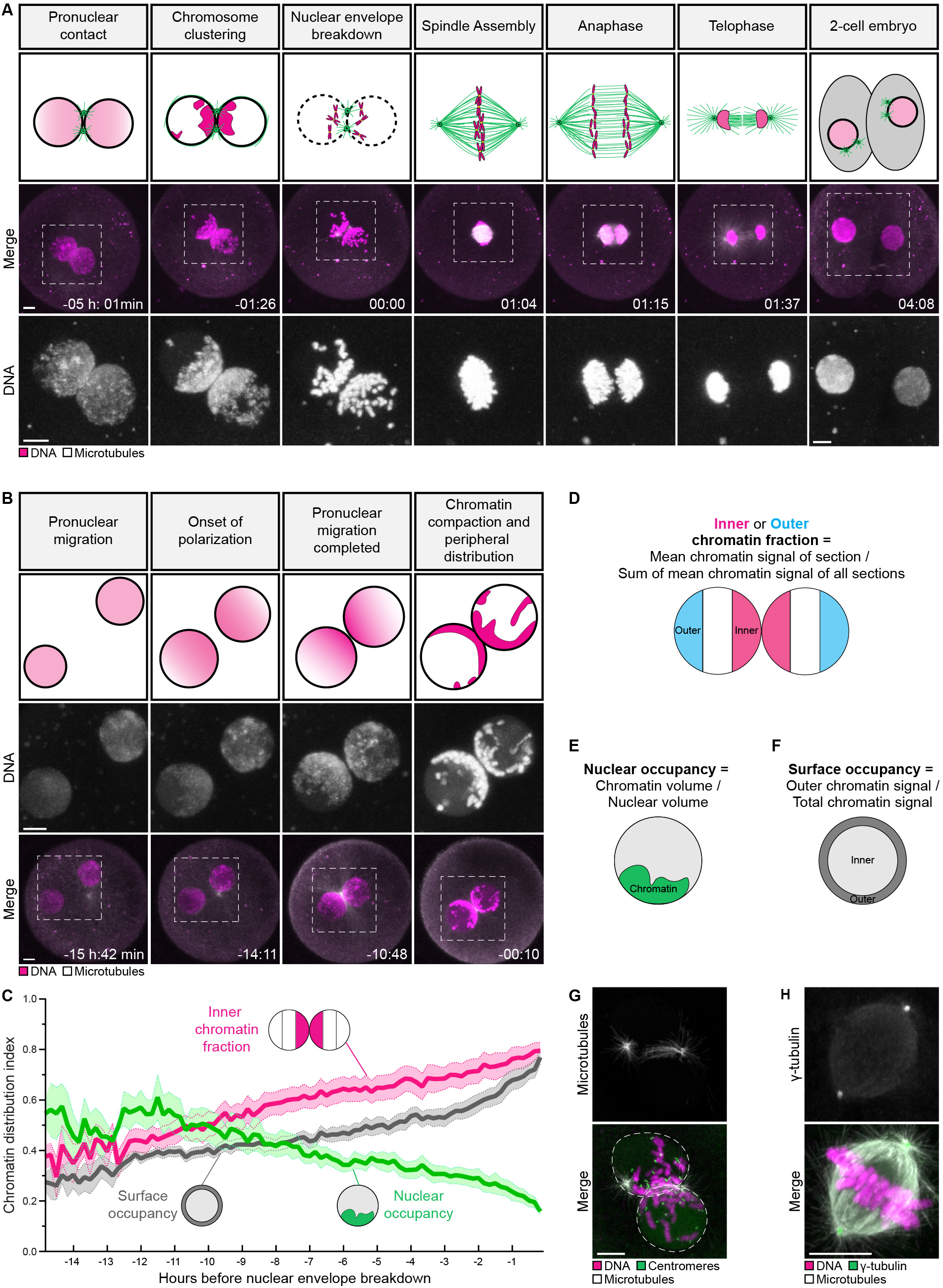
Parental genome unification initiates in intact nuclei upon fertilization. (A) (Top) Schematic of chromatin organization and spindle assembly in zygotes. Magenta, chromatin. Green, microtubules. (Bottom) Representative stills from time-lapse movies of zygotes. Gray, microtubules (mClover3-MAP4-MTBD). Magenta, DNA (H2B-mScarlet). Outlined regions are magnified below. Time, hours:minutes, 00:00 is nuclear envelope breakdown. Z-projections, 11 sections every 2.50 µm. (B) (Top) Schematic of chromatin organization during pronuclear migration. (Bottom) Representative stills from time-lapse movies of pronuclear migration. Gray, microtubules (mClover3-MAP4-MTBD). Magenta, DNA (H2B-mScarlet). Outlined regions magnified above. Time, hours:minutes, 00:00 is nuclear envelope breakdown. Z-projections, 8 sections every 2.50 µm. (C) Quantification of chromatin distribution within pronuclei using the inner chromatin fraction index (magenta), nuclear occupancy index (green), and surface occupancy index (gray). Solid lines represent means of ten pronuclei belonging to five zygotes obtained from three independent experiments. Shaded areas represent standard error of the mean. (D-F) Schematics of the chromatin distribution indices used in (C). (G) Representative immunofluorescence images of a zygote upon nuclear envelope breakdown. Gray, microtubules (α-tubulin). Magenta, DNA (DAPI). Green, centromeres (ACA). (H) Representative immunofluorescence images of a zygotic spindle. Gray, microtubules (α-tubulin). Magenta, DNA (DAPI). Green, γ-tubulin. Scale bars, 10 µm.

Chromatin clustering began in the early stages of pronuclear migration, and accelerated as the chromosomes condensed during the final 3-4 hours before nuclear envelope breakdown (Figure 1C). As a consequence of chromosome clustering and condensation, the chromatin occupied less than 20% of the nucleus just before nuclear envelope breakdown (Figures 1C, 1E, and S1C). Before nuclear envelope breakdown, all chromosomes were associated with the periphery of the pronuclei, including the few chromosomes located away from the pronuclear interface (Figures 1A-1C, 1F, and S1D). In intact somatic nuclei and pronuclei of other systems, the chromatin remains uniformly distributed and much less condensed prior to cell division (Courtois et al., 2012; Gonczy et al., 1999; Magidson et al., 2011). Thus, the clustering of parental genomes toward each other within intact pronuclei (hereafter referred to as “pre-unification”) and the early chromosome condensation were completely unexpected.

Upon nuclear envelope breakdown, the chromosomes were captured by microtubules emanating from the two centrosomes. The centrosomes served as major microtubule organizing centers (Figure 1G; Movie S1), and rapidly assembled a spindle with two focused poles (Figure 1H; Movie S1). Interestingly, the maternal and paternal chromosomes continued to occupy partially distinct territories upon nuclear envelope breakdown and on the first mitotic spindle (Figure S1E; Movie S2), consistent with work in *C. elegans*, mouse, and humans (Bolkova and Lanctot, 2016; Mayer et al., 2000; Reichmann et al., 2018; van de Werken et al., 2014). Subsequently, the zygote progressed into anaphase and divided into two equal daughter cells, each with a parental genome complement (Figure 1A).

### Parental genome pre-unification ensures accurate chromosome segregation

We reasoned that pre-unification of the parental genomes may facilitate their capture upon nuclear envelope breakdown. The two pronuclei spanned a width of ∼ 47 µm just before nuclear envelope breakdown (Figure S1F), which is much larger than the width of the somatic nucleus in a typical mitotic cell (Milo and Phillips, 2016), and much longer than mitotic microtubules in bovine zygotes (8.7±2.6 µm) (Figure S1G). These data suggest that chromosome capture might be inefficient if the chromosomes were uniformly distributed throughout the two pronuclei. Pre-unification of the parental genomes at the pronuclear interface reduces the total chromosome volume, and thus could promote efficient chromosome capture by microtubules.

To test this hypothesis, we took advantage of our observation that some zygotes failed to cluster their chromatin in one of the two pronuclei. Specifically, we observed two major classes of clustering defects. The first class is characterized by less condensed, unclustered chromatin in one pronucleus (Figure 2A), which we termed uncondensed. The second class is characterized by peripherally localized and condensed chromatin that is not at the interface of the two pronuclei, which we termed unclustered (Figure 2A).

**Figure 2.**
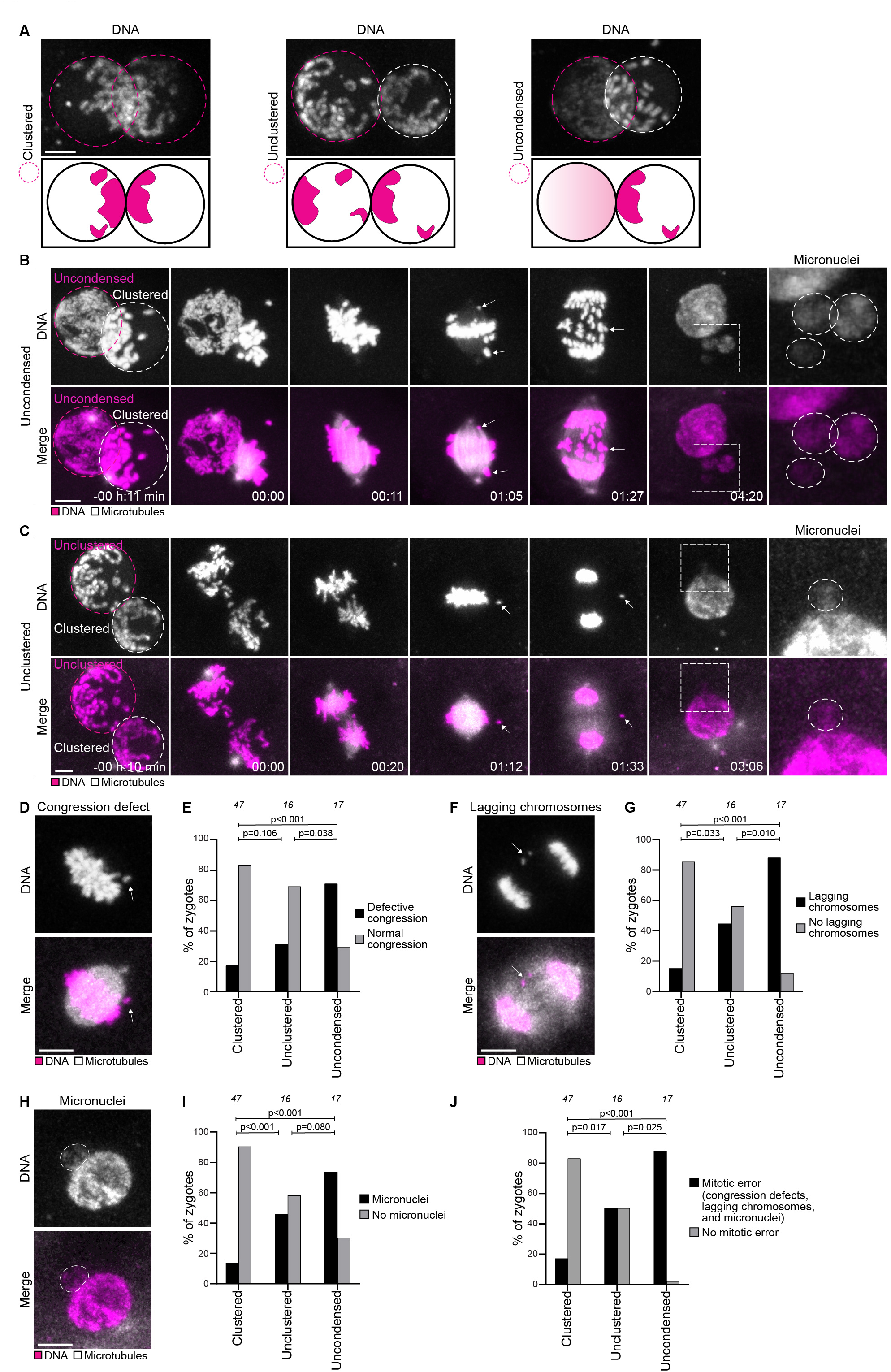
Parental genome pre-unification ensures accurate chromosomes segregation. (A) Representative stills from time-lapse movies (top) and schematics (bottom) of zygotes pronuclei before nuclear envelope breakdown classified as clustered (non-defective, left); unclustered (middle); uncondensed (right). Gray, DNA (H2B-mScarlet). Magenta dashed line indicates the pronucleus determining the specific category. Z-projections, 10 sections every 2.50 µm. (B-C) Representative stills from time-lapse movies of bovine zygotes before and after nuclear envelope breakdown. (B) shows a zygote classified as uncondensed. (C) shows a zygote classified as unclustered. Gray, microtubules (mClover3-MAP4-MTBD). Magenta, DNA (H2B-mScarlet). Arrows indicate misaligned and lagging chromosomes that form micronuclei, as highlighted by dashed box (magnification on the right). Time, hours:minutes, 00:00 is nuclear envelope breakdown. Z-projections, 12 sections every 2.50 µm. (D-J) Representative images and frequencies of zygotes in indicated groups having defective chromosome congression (D, E), lagging chromosomes (F, G), micronuclei in 2-cell embryos (H, I), and abnormal mitosis (J). Gray, microtubules (mClover3-MAP4-MTBD). Magenta, DNA (H2B-mScarlet). Z-projections, respectively 5, 4, and 5 sections every 2.50 µm. Data in (E, G, I, J) are from eleven independent experiments. The number of analyzed zygotes is specified in italics. Indicated *p*-values calculated using Fisher’s exact tests. Scale bars, 10 µm.

We manually identified zygotes with an uncondensed pronucleus based on their chromatin configuration in the frame before nuclear envelope breakdown (Figure 2A), and discerned zygotes with an unclustered pronucleus from zygotes with clustered pronuclei using algorithms that quantify chromatin distribution (see Methods) (Figures 1D-1F and S2A-S2D). These three groups of zygotes were followed as they progressed through the first mitotic division (Figures 1A, 2B, 2C, S3A, and S3B; Movies S1, S3, and S4) and scored for phenotypes associated with defective chromosome capture, including: (1) compromised congression on the metaphase plate (Figures 2D and 2E), (2) lagging chromosomes during anaphase (Figures 2F and 2G), and (3) formation of micronuclei (Figures 2H and 2I).

By these criteria, the vast majority of zygotes with an uncondensed pronucleus showed serious defects during mitosis (Figure 2J). Chromosomes from the uncondensed pronucleus showed delays in both association with the spindle and congression on the metaphase plate (Figures 2B and S3A; Movie S3). Indeed, 71% of uncondensed zygotes failed to congress all their chromosomes on the metaphase plate by the time of anaphase onset (Figure 2E), and anaphase onset was significantly delayed (Figure S2E). Additionally, 88% of these zygotes had multiple lagging chromosomes during anaphase (Figure 2G). To investigate which pronucleus gives rise to the lagging chromosomes, we selectively photobleached the chromosomes in the peripheral clustered pronucleus thus leaving the chromosomes in the uncondensed pronucleus unbleached and distinguishable (Figure S4A; Movie S3). We found that 89% of lagging chromosomes originated from the uncondensed pronucleus (Figures S4B and S4C; Movie S3). These data are consistent with defective capture of chromosomes originating from the uncondensed pronucleus.

Zygotes with an unclustered pronucleus also showed prominent defects in mitosis. Chromosomes in the distal regions of the pronuclei often remained separated from the main chromosome cluster upon nuclear envelope breakdown, and the time to congress chromosomes on the metaphase plate was delayed (Figures S2F and S3B; Movie S4). Although most zygotes in this class eventually succeeded in congressing their chromosomes on the metaphase plate (Figure 2E), 50% of them had lagging chromosomes during anaphase, indicating incorrect attachment to microtubules (Figure 2G). Interestingly, anaphase onset was not significantly delayed (Figure S2E), suggesting that zygotes can progress into anaphase without delay despite incorrect kinetochore-microtubule attachments. These data suggest that abnormal kinetochore microtubule attachments are not detected by zygotes, and/or do not prevent progression into anaphase. Rapid chromosome capture upon nuclear envelope breakdown may hence be of particular importance in zygotes, in order to provide sufficient time to correct erroneous attachments before anaphase onset.

To test if the unclustered chromosomes were more likely to lag in these zygotes, we selectively photobleached the clustered chromosomes at the pronuclear interface, leaving the distal chromosomes unbleached (Figure S4D; Movie S4). Consistent with our hypothesis, 69% of detected lagging chromosomes were unbleached, indicating that inside the pronuclei they were distal from the pronuclear interface (Figure S4E). Bleaching did not harm the zygotes, because most bleached zygotes lacked chromosome segregation defects, similar to untreated zygotes (Figure S4F). Together, these data indicate that distal chromosomes are more likely to lag during anaphase than clustered chromosomes.

Lagging chromosomes were generally more frequent in zygotic mitosis than in mitosis of somatic cells (Thompson and Compton, 2008, 2011), suggesting an increased rate of incorrect kinetochore-microtubule attachments or a less efficient error correction mechanism. The lagging chromosomes were often subsequently encapsulated in micronuclei (Figures 2H and 2I). Most embryos with micronuclei had both chromosome congression defects and lagging chromosomes (Figure S2G). Thus, defective chromosome congression on the first mitotic spindle is a major factor contributing to the frequent appearance of lagging chromosomes and, in turn, micronuclei in two-cell embryos.

In total, 90% of zygotes with an uncondensed pronucleus, and 50% of zygotes with an unclustered pronucleus had defects during the first mitotic division (Figure 2J). This is in stark contrast to zygotes with two clustered pronuclei: only 20% of these had defects (Figure 2J). Together, these data establish that parental genome pre-unification is required to accurately segregate chromosomes and prevent the formation of micronuclei (Figure 7A). Chromosome segregation errors in the zygote will give rise to aneuploid 2-cell embryos, precluding the development of a healthy embryo (Kort et al., 2016; Vazquez-Diez et al., 2016).

### Centrosome positions determine sites of chromosome clustering and accuracy of chromosome segregation

Having established the functional importance of parental genome pre-unification, we wanted to investigate the underlying mechanism. We noticed that the chromosomes normally moved to the position of the duplicated centrosomes (Figure 1A; Movie S1). In the vast majority of zygotes, one or two centrosomes were located at the interface between the two pronuclei (Figure 3A) and were always associated with chromosomes (Figures 3B and 3C). One of the two centrosomes localized sometimes distal from the pronuclear interface, but was also associated with chromosomes in 98% of all cases (Figures 3B and 3C). Interestingly, zygotes with centrosomes positioned away from the pronuclear interface failed to pre-unite the parental genomes (Figures 3D and 3E) and were more likely to have chromosome segregation errors (Figure 3F). After nuclear envelope breakdown, the parental genomes of zygotes with centrosomes positioned away from the pronuclear interface remained separate for a longer time period and only united with a delay (Figure 3G).

**Figure 3.**
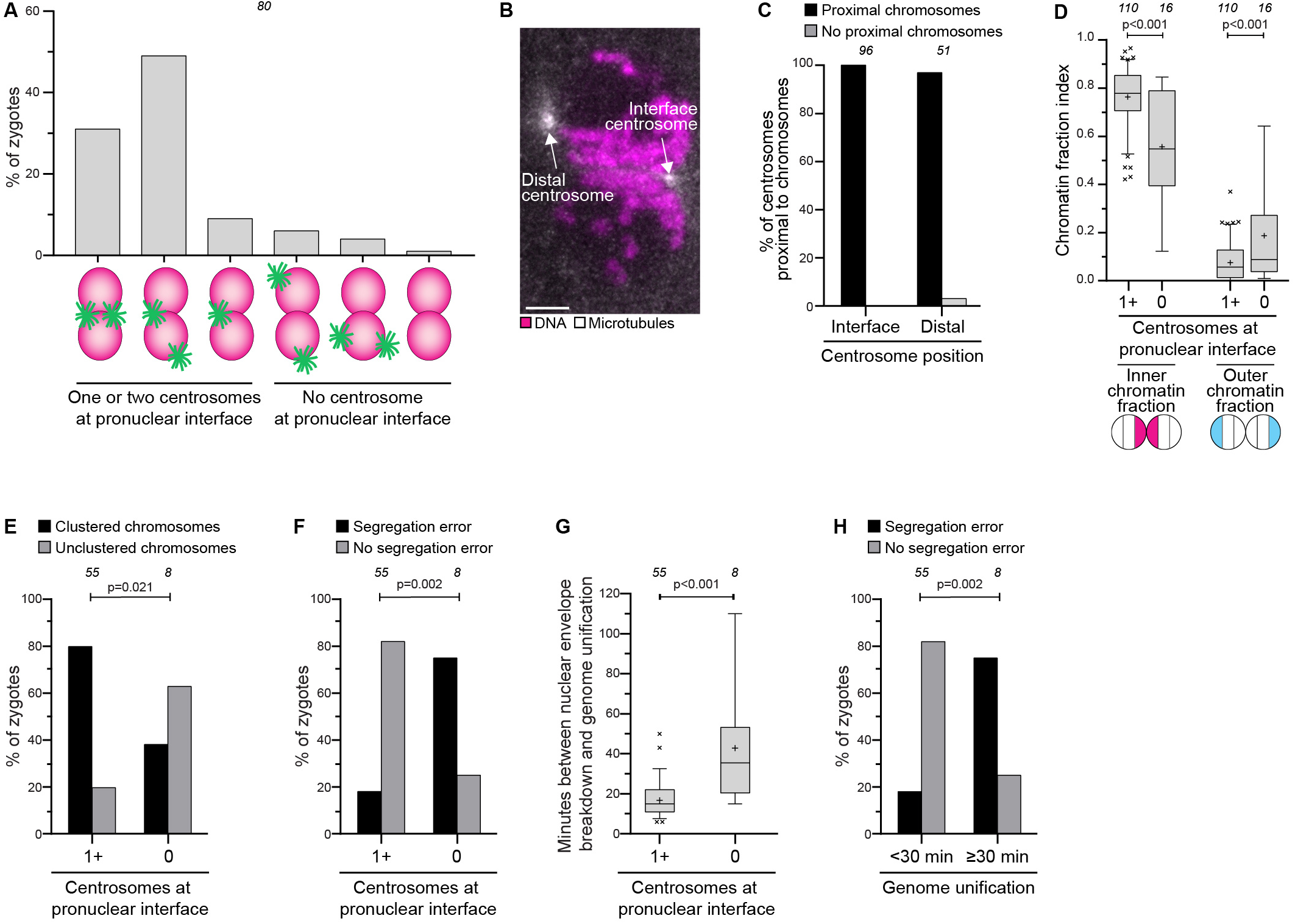
Centrosome positions determine sites of chromosome clustering and accuracy of chromosome segregation. (A) Centrosome localization before nuclear envelope breakdown in the indicated configurations. (B) Representative still from a time-lapse movie of a zygote before nuclear envelope breakdown. Gray, microtubules (mClover3-MAP4-MTBD). Magenta, DNA (H2B-mScarlet). Arrows specify the distal and the interface centrosomes. (C) Centrosomes located away from the pronuclear interface that are proximal or not to chromosomes in pronuclei. (D) Inner and outer chromatin fraction indices in indicated groups. (E) Zygotes with clustered or unclustered chromosomes before nuclear envelope breakdown with different centrosome positioning as indicated. (F) Zygotes having chromosome segregation errors during mitosis with different centrosome positioning as indicated. (G) Time between nuclear envelope breakdown and the unification of the parental genomes in zygotes with different centrosome positioning as indicated. (H) Zygotes having chromosome segregation errors during mitosis where genome unification take place within 30 minutes after nuclear envelope breakdown or later as indicated. Zygotes having a pronucleus with uncondensed chromatin at nuclear envelope breakdown have been excluded from the quantifications in C, D, E, F, G, and H to avoid accounting for the role of incomplete chromosome condensation at nuclear envelope breakdown. Data are from eleven independent experiments. The number of analyzed zygotes (A, E, F, G, H), centrosomes (C), and pronuclei (D) is specified in italics. Indicated *p*-value calculated using unpaired two-tailed Student’s t test (D, G) and Fisher’s exact test (E, F, H). Scale bar, 10 µm.

Zygotes with a delay in genome unification were also generally more likely to have mitotic errors (Figure 3H), emphasizing the importance of rapidly unifying the parental genomes upon nuclear envelope breakdown. Together, these data suggest that centrosomes determine the site of chromosome clustering and that their presence at the pronuclear interface promotes the rapid and error-free unification of the parental genomes.

### Centrosomes, microtubules, and dynein pre-unite chromosomes within intact pronuclei

Our hypothesis that centrosomes drive chromosome clustering predicts that detaching centrosomes from pronuclei would lead to defects in chromosome clustering. Work in *C. elegans* and zebrafish zygotes has established that centrosomes are coupled to nuclei through the LINC complex and dynein (Bone and Starr, 2016; Lindeman and Pelegri, 2012; Malone et al., 2003). This coupling requires the KASH5 subunit of the LINC complex, which can be blocked with a KASH5 dominant-negative (KASH5-DN) fragment (Stewart-Hutchinson et al., 2008). We expressed a KASH5-DN in bovine zygotes (Figure S5A), and observed the displacement of centrosomes from the nuclear envelope, failure in pronuclear migration, and, eventually, assembly of two separate spindles (Figures S5B-S5G; Movie S5). These data suggest that centrosome coupling to the nuclear envelope in bovine zygotes also relies on the LINC complex and is necessary for pronuclear migration, consistent with work in non-mammalian embryos (Cowan and Hyman, 2004; Wuhr et al., 2010).

KASH5-DN zygotes displayed significantly decreased chromosome clustering, consistent with the notion that centrosomes drive chromosome clustering (Figures 3A and 3B; Movie S5). However, recruitment of chromosomes to the nuclear envelope was not affected (Figure S5H).

Next, we investigated how centrosomes direct chromosome movements. Centrosomal microtubules directly capture chromosomes via their kinetochores during somatic cell mitosis (Heald and Khodjakov, 2015). However, in zygotes, microtubules did not permeate the nuclear envelope during chromosome clustering, but instead wrapped around the pronuclear envelopes (Figure 4D). Moreover, centromeres were located in the inner region of the pronuclei, away from the nuclear envelope (Figures S6A-SC), suggesting that centrosomes cluster the chromosomes via a different mechanism.

**Figure 4.**
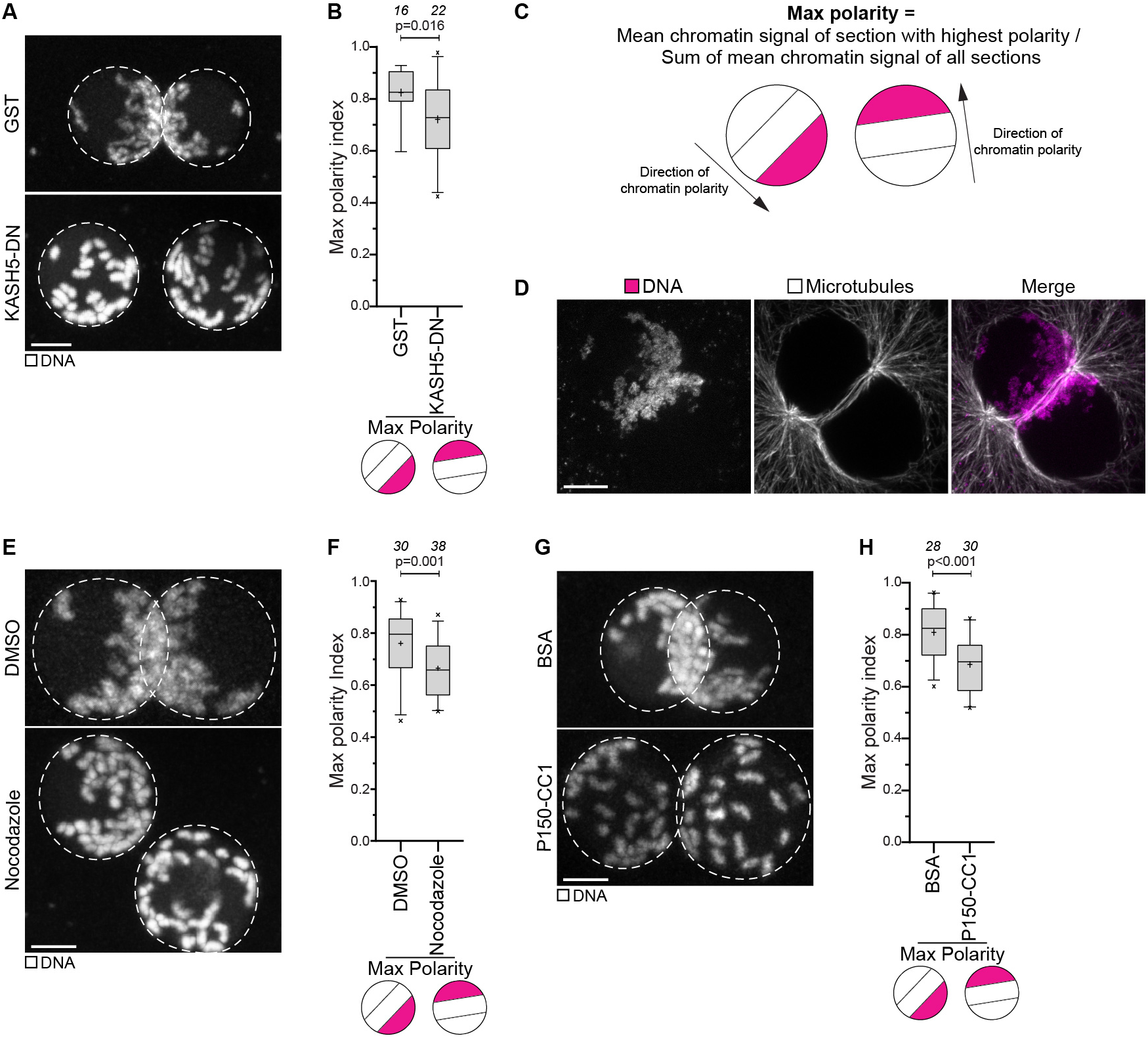
Centrosomes, microtubules, and dynein pre-unite chromosomes within intact pronuclei. (A) Representative images of zygotes before nuclear envelope breakdown expressing GST or KASH5-DN. Gray, DNA (H2B-mScarlet). Dashed lines mark pronuclei. Z-projections, 9 sections every 3.08 µm. (B) Max polarity index (see (C)) in zygotes expressing GST or KASH5-DN. (C) Schematics of the max polarity index. (D) Representative immunofluorescence images of a zygote with microtubules wrapped around the pronuclei. Gray, microtubules (α-tubulin). Magenta, DNA (DAPI). Z-projection, 35 sections every 0.1 µm. (E-H) Representative images and max polarity indices in zygotes treated with DMSO or nocodazole (E, F) or injected with BSA or P150-CC1 (G, H). The last time point before nuclear envelope breakdown is shown. Gray, DNA (H2B-mScarlet). Dashed lines mark pronuclei. Z-projections, respectively 8 and 12 sections every 2.50 µm. Data are from four (B, H) or six (F) independent experiments. The number of analyzed pronuclei is specified in italics. Indicated *p*-values calculated using unpaired two-tailed Student’s t-test. Scale bars, 10 µm.

The perinuclear microtubules that wrapped around the nuclear envelope appeared to be in a suitable arrangement for mediating chromosome transport towards centrosomes, as they covered most of the pronuclear surfaces and were highly enriched at the pronuclear interface, where at least one of the centrosomes was located (Figure 4D). Adapters in the nuclear envelope can link chromosomes to microtubules and thereby transduce forces across the nuclear envelope (Zeng et al., 2018). Transport towards centrosomes in interphase is typically mediated by the minus-end directed motor protein dynein (Malone et al., 2003; Quintyne et al., 1999). Thus, we hypothesized that dynein might transport the chromosomes to the centrosome via adaptors that bridge the nuclear envelope.

To test this hypothesis, we first investigated if microtubules are required for chromosome clustering by treating zygotes with the microtubule-depolymerizing drug nocodazole. In support of transport along microtubules, nocodazole not only blocked pronuclear migration (Figure S5I; Movie S5), but significantly reduced the clustering of chromosomes within pronuclei (Figures 4E and 4F). Consistent with the effects of KASH5-DN, chromosomes still relocated to the nuclear envelope, suggesting that the recruitment of chromosomes into the nuclear periphery is independent of microtubules and centrosomes (Figure S5J).

To test for an involvement of dynein, we purified and injected the C-terminus of the dynein interaction partner dynactin (P150-CC1), which blocks the activity of the dynein-dynactin complex (Quintyne et al., 1999). P150-CC1 led to a significant reduction of chromosome clustering at the pronuclear interface (Figures 4G, 4H, and S5KL; Movie S5). Overall, our data suggest that dynein mediates the transport of chromosomes along microtubules towards the centrosomes.

### Nuclear pore complexes cluster with chromatin at the pronuclear interface

The dynein-dependent movement of chromosomes toward the centrosome in zygotes is reminiscent of the mechanisms involved in forming telomere bouquets established in the early stages of meiosis (Zeng et al., 2018). The telomeres cluster in proximity to the centrosome in preparation for meiotic recombination (Sato et al., 2009; Shibuya et al., 2014). However, in stark contrast to meiotic cells, telomeres were located in the central region of zygotic pronuclei, and were often enriched at nucleoli together with centromeres (Figures S6D-S6F), consistent with reports in mouse and human zygotes (Jachowicz et al., 2013; van de Werken et al., 2014). Moreover, the inner nuclear membrane protein SUN1 was distributed along both pronuclear envelopes, without specific enrichment on peripheral chromosomes (Figure S6G). This localization is in contrast to the clustered appearance of SUN1 in proximity to the centrosome in meiotic cells, but it is consistent with observations in *C. elegans* zygotes (Minn et al., 2009). These data suggest that dynein moves chromosomes in zygotes by a mechanism that is distinct from telomere movement in early meiosis.

Studies in somatic cells have shown that dynein can bind to nuclear pore complexes (Bolhy et al., 2011; Splinter et al., 2010) and that this interaction is required for nuclear migration in brain progenitor cells (Hu et al., 2013). Interestingly, previous work reported that chromosomes associate with nuclear pore complexes in bovine embryos from the 2-cell stage onwards (Popken et al., 2015). This led us to investigate if dynein pulls the chromosomes to the centrosome via nuclear pore complexes. Transport of nuclear pore complexes toward centrosomes was indeed evident in videos of live zygotes labelled with the nucleoporin Pom121 (Figure 5A; Movie S6). Although individual pores could not be resolved on intact nuclei, annulate lamellae, membrane stacks enriched in nuclear pore complexes, were prominently recruited toward centrosomes as the two pronuclei moved inwards (Figure 5A; Movie S6, arrowheads).

**Figure 5.**
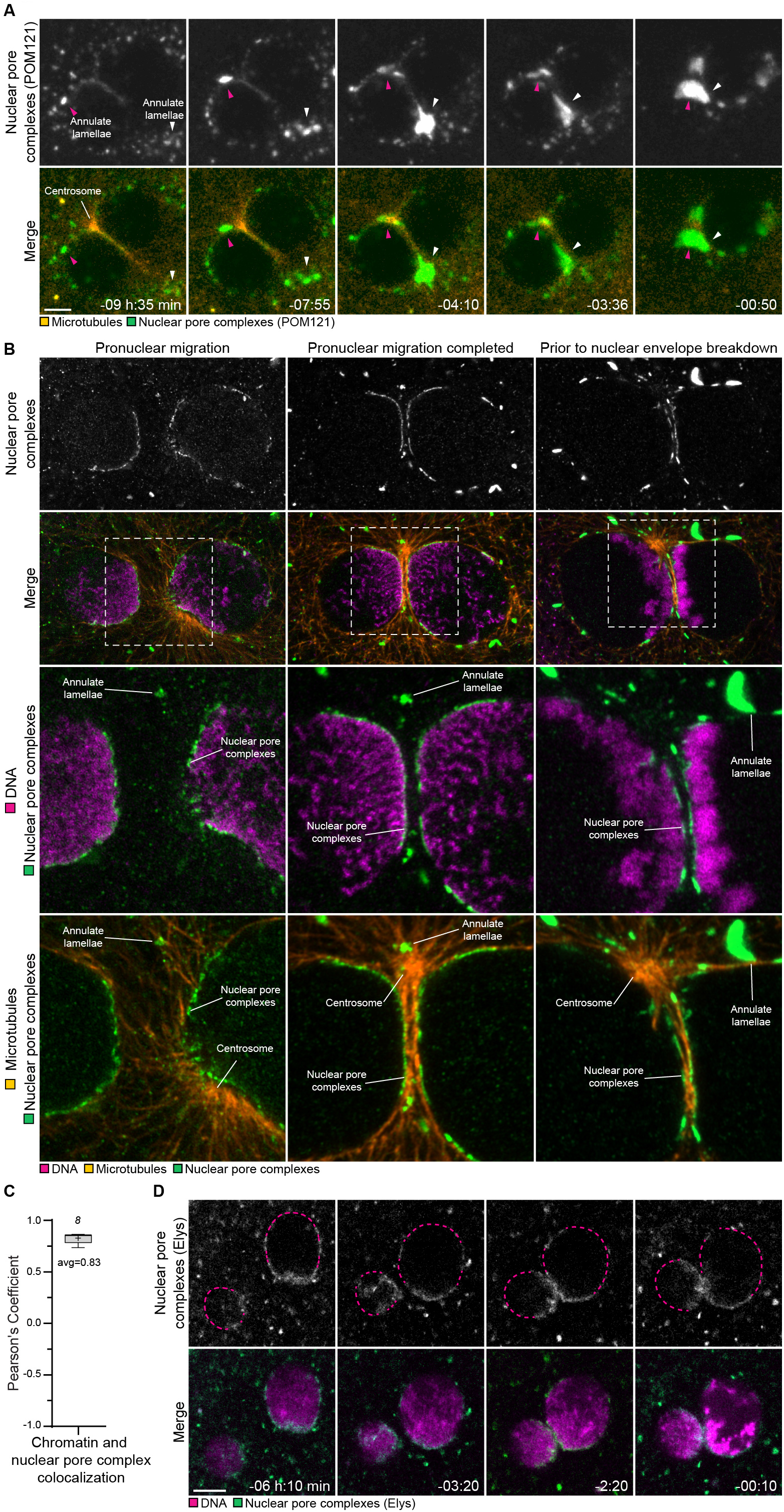
Nuclear pore complexes cluster with chromatin at the pronuclear interface. (A) Representative stills from time-lapse movies of zygotes expressing mClover3-MAP4-MTBD (microtubules, orange) and POM121-mScarlet (nuclear pores complexes, green). Magenta and white arrowheads indicate two patches of annulate lamellae clustering toward centrosomes and pronuclear interface. Time, hours:minutes, 00:00 is nuclear envelope breakdown. Single confocal microscopy sections. (B) Representative immunofluorescence images of zygotes before pronuclear apposition (left), before chromosome condensation (middle), and after chromosome condensation (right). Orange, microtubules (respectively, β-tubulin, α-tubulin, and α-tubulin). Magenta, DNA (DAPI). Green, nuclear pore complexes (respectively Nup98, NPC-Mab414, and NPC-Mab414). Outlined regions magnified on the bottom. Centrosomes, nuclear pore complexes, and annulate lamellae are indicated. Single sections Airyscan microscopy. (C) Pearson’s coefficient quantifying the co-localization at the nuclear envelope of nuclear pore complex and chromatin signals. +1 indicates perfect co-localization, −1 indicates exclusion. (D) Representative stills from time-lapse movies of zygotes expressing bElys-mClover3 (nuclear pore complexes, green) and H2B-mScarlet (DNA, magenta). Dashed line indicates region of the nuclear envelope devoid of nuclear pore complexes. Time, hours:minutes, 00:00 is nuclear envelope breakdown. Single confocal microscopy sections, except for first image where two sections were Z-projected to visualize both pronuclei. Data in (C) are from two independent experiments. The number of analyzed zygotes is specified in italics. Scale bars, 10 µm.

To gain better resolution, we performed Airyscan super-resolution microscopy of nuclear pore complexes in fixed zygotes. We found that nuclear pore complexes were unevenly distributed along the nuclear envelope, clustered at the pronuclear interface and in direct proximity to the clustered chromosomes (Figure 5B; Movie S7). We quantified this co-localization, obtaining a Pearson’s correlation coefficient of 0.83 (Figure 5C). In contrast, Lamin A/C and Lamin B1 were distributed along the entire nuclear envelope (Figures S6H and 6I).

The clusters of nuclear pore complexes were closely associated with microtubules running along the interface between the two pronuclei (Figures 5B; Movie S7), consistent with dynein pulling chromosomes along perinuclear microtubules via nuclear pore complexes.

Live imaging further demonstrated that the polarization of nuclear pore complexes is established in parallel to chromatin clustering as the pronuclei move toward each other (Figure 5D). Altogether these data are consistent with a model whereby chromosome clustering is driven by dynein-dependent pulling on nuclear pore complexes (Figure 7B).

### Nucleoli as a read-out of chromosome clustering and blastocyst development in human embryos

We next set out to investigate if chromosome pre-unification also occurs in human zygotes. Labelling of chromosomes in human embryos is not permitted as part of fertility treatments, and fertilizing human eggs for research is forbidden in many countries, including Germany. However, in vitro fertilization clinics routinely image human embryos by transmission microscopy (Cruz et al., 2011). These videos are typically recorded at sufficient resolution to detect the position of the nucleolar precursor bodies (hereafter, nucleoli), which are associated with centromeres and telomeres in bovine and in human zygotes (Figures S6A and S6D) (van de Werken et al., 2014). By imaging nucleoli in live bovine zygotes, we confirmed their co-localization with chromatin (Figure S6J), indicating that nucleoli can serve as proxy for chromatin distribution in zygotes.

Interestingly, previous studies had suggested that the distribution of the nucleoli can predict the developmental efficiency of human embryos (Tesarik and Greco, 1999). However, these studies were carried out prior to routine live microscopy of embryos, and different studies had come to varying conclusions, possibly due to the evaluation of single time points (Fulka et al., 2015; Guerif et al., 2007; James et al., 2006; Swain, 2013). More recent studies that took advantage of live microscopy combined multiple parameters and thus no clear result emerged for clustering of nucleoli (Azzarello et al., 2012; Coticchio et al., 2018; Faramarzi et al., 2018). We hence repeated this analysis by scoring videos of live human embryos for clustered and non-clustered nucleoli (Figures 6A and 6B; Movie S8).

**Figure 6.**
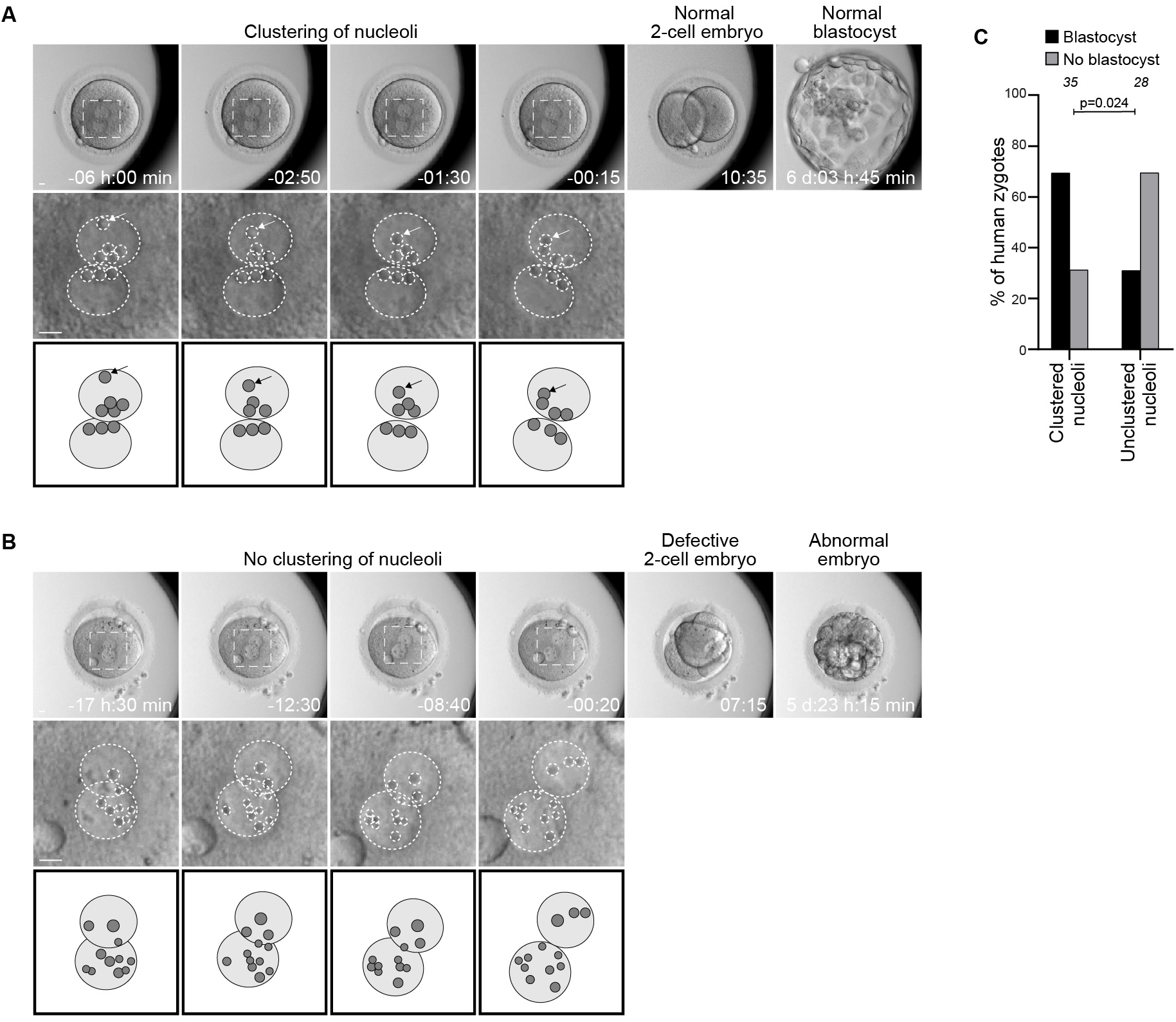
Nucleoli as a read-out of chromosome clustering and blastocyst development in human embryos. (A-B) (Top) Representative stills from time-lapse movies of a human zygote that develops (A) or fails to develop (B) into a blastocyst. Zygotes have clustered (A) or unclustered (B) nucleoli at the pronuclear interface. (Middle) Magnifications of the regions outlined above. Dashed lines indicate nucleoli. (Bottom) Schematics of the pronuclei and nucleolar distribution. Arrows indicate a nucleolus that moves toward the pronuclear interface. (C) Human zygotes with clustered or unclustered nucleoli that develop into blastocyst or develop abnormally in indicated groups. The number of analyzed embryos is specified in italics. Indicated *p*-value calculated using Fisher’s exact test. Scale bars, 10 µm.

**Figure 7.**
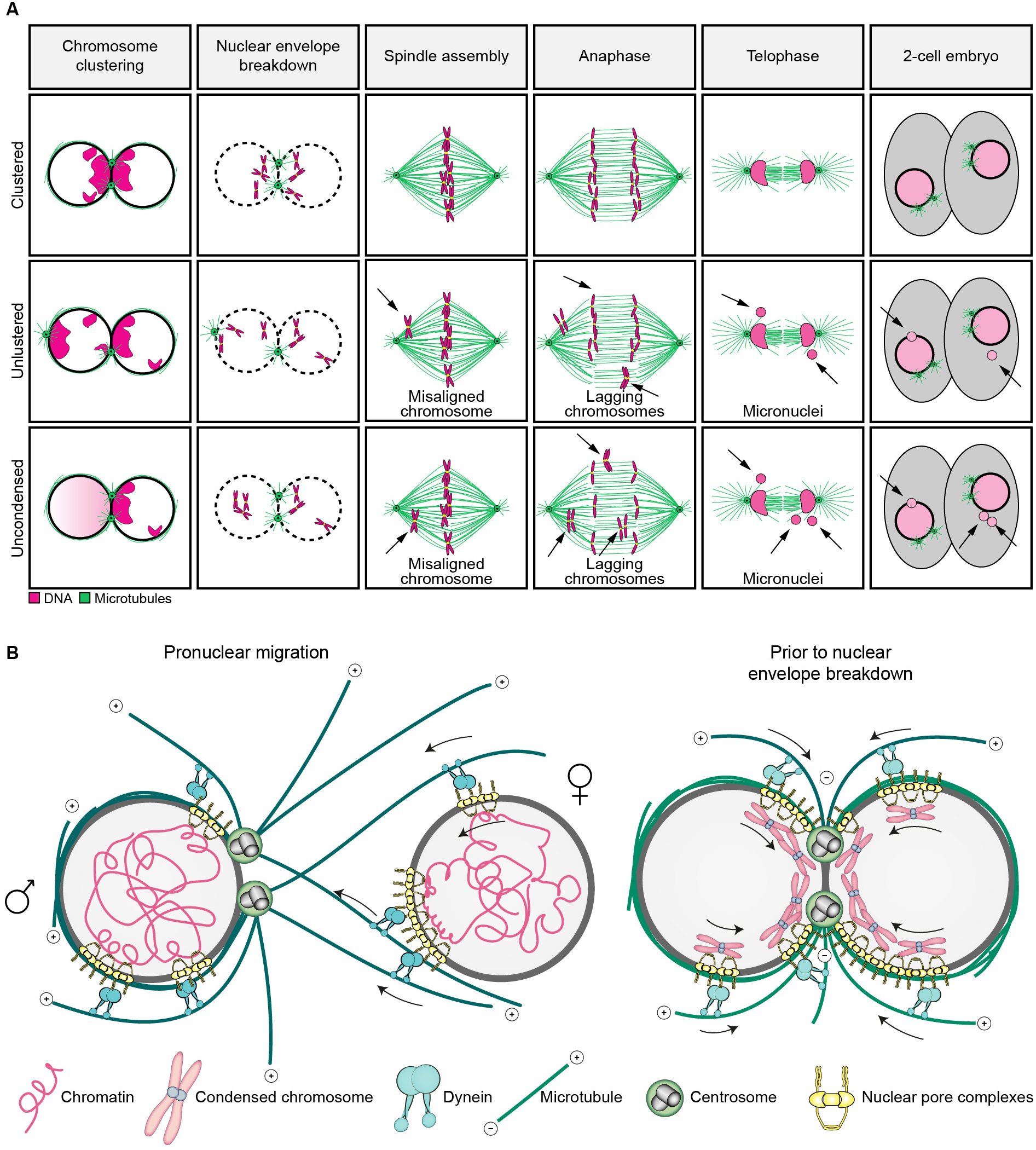
Models illustrating the mechanism of chromosome clustering and parental genomes pre-unification. (A) Model for the mechanism of chromatin clustering at the pronuclear interface before (left) and after (right) migration completion. Gray, nucleoplasm. Magenta, chromatin and chromosomes. Yellow, nuclear pores. Cyan, dynein. Green, microtubules and centrosomes. Microtubule polarity is indicated by + and -. Arrows indicated dynein directionality. Female and male pronuclei are indicated by ♀ and ♂, respectively. (B) Schematics of mitosis in zygotes having clustered (top), unclustered (middle), or uncondensed (bottom) chromosomes. Depending on the chromatin organization before nuclear envelope breakdown mitotic errors occur. Magenta, chromatin. Green, microtubules. Arrows point to defects causing chromosome segregation errors, such as misaligned and lagging chromosomes, and to micronuclei.

We found that 56% of human zygotes had clustered nucleoli at the pronuclear interface (Figure 6C), which is strikingly similar to the 59% of bovine zygotes with clustered chromatin that we observed. This configuration is likely to correspond to a polarized chromatin distribution, which was observed in an early study of human zygotes but has not been further investigated (Van Blerkom et al., 1995). In some cases, we could also observe nucleoli moving within the pronuclei towards the pronuclear interface region (Figure 6A, arrow), suggesting active clustering similar to what we observed for chromatin in bovine zygotes. Strikingly, human zygotes with clustered nucleoli were significantly more likely to develop into blastocysts than zygotes with scattered nucleoli (Figure 5C; Movie S8). These data suggest that the clustering of nucleoli at the pronuclear interface has a positive impact on human embryo development, consistent with our observations that chromatin clustering promotes proper development of bovine embryos.

## Discussion

Here, we established a high-resolution live cell microscopy system to study the development of bovine embryos as a model for human embryos. Using this system, we discovered an unexpected, error-prone mechanism that unites the maternal and paternal chromosomes. In particular, the parental genomes cluster in close proximity to each other at the interface between their intact pronuclei, a process that we defined as parental genome pre-unification. Our data suggest that clustering is achieved by dynein, which transports chromosomes along perinuclear microtubules toward the centrosomes as the pronuclei migrate inwards (Figure 7B). Parental genome pre-unification increases the efficiency of chromosome capture by the newly assembling spindle, and thus prevents chromosome segregation errors (Figure 7A).

Pronuclear migration and chromosome clustering are not only established simultaneously, but also the molecular players driving both processes – microtubules, dynein, nuclear pore complexes, and centrosomes – are intriguingly similar (Hu et al., 2013; Lindeman and Pelegri, 2012; Malone et al., 2003; Payne et al., 2003). Our data are consistent with a unified model whereby pronuclear migration and chromatin clustering are two tightly interwoven processes, simultaneously established by the same cellular machinery, with the common aim of uniting the parental genomes. In this model, dynein associates with nuclear pore complexes and transports the pronuclei toward each other along centrosome-nucleated microtubules. Pulling via nuclear pore complexes not only brings the two pronuclei into close proximity, but it simultaneously pre-unites the parental genomes at the pronuclear interface to facilitate their rapid capture and union on the first mitotic spindle (Figure 7).

Bovine and human embryos are frequently aneuploid, with multiple errors resulting from defects during the first cell divisions (Destouni et al., 2016; Lee and Kiessling, 2017; McCoy, 2017; McCoy et al., 2015b; Vera-Rodriguez et al., 2015). Defects during parental genome unification in the zygote are likely to be a major cause of high aneuploidy rates in embryos and abnormal embryo development. Interestingly, a high number of zygotes displayed clustering defects and a delay in parental genome unification, which were both highly correlated with lagging chromosomes, a frequent cause of aneuploidy. The pre-unification of the parental chromosomes is hence a particularly critical and sensitive step in embryo development.

Our data also reveal fundamental differences in the way that zygotes and normal somatic cells progress through mitosis. Apart from their large size and the presence of two nuclei, zygotes also display a high degree of chromatin polarization toward the pronuclear interface, condensed chromosomes at the nuclear periphery before nuclear envelope breakdown, and a highly asymmetric distribution of nuclear pore complexes.

Chromosome condensation and recruitment into the nuclear periphery may help to expose kinetochores and bring them into close proximity with the centrosomes to facilitate their capture. Kinetochores buried within an uncondensed chromatin mass, such as those in uncondensed pronuclei, make microtubule contacts later and often stay incorrectly attached, as evident from a large number of lagging chromosomes in this group. The fact that 21% of zygotes have an uncondensed nucleus before nuclear envelope breakdown implies that asynchronous chromosome condensation in the two pronuclei is a frequent phenomenon in bovine zygotes and a major cause of errors during the first mitotic division. This number is strikingly similar to the 25% of human cleavage embryos displaying more than 3 whole chromosomes aneuploidy (McCoy et al., 2015b), also referred to as chaotic aneuploidy (McCoy, 2017).

Chromosome clustering might occur specifically in zygotes because their chromosomes are spread over a much larger volume than in somatic mitotic cells, and hence are more difficult to capture and unite by two centrosomes. The position of the centrosomes plays a crucial role in this process, as zygotes with centrosomes positioned away from the pronuclear interface fail to cluster the chromosomes, unify the parental genomes later, and are more likely to show chromosome segregation errors.

Interestingly, mouse zygotes have multiple acentriolar microtubule organizing centers distributed on the surface of the two pronuclei, which may facilitate the rapid capture of chromosomes upon nuclear envelope breakdown (Courtois et al., 2012). This alternative and potentially more efficient capture mechanism may explain why mice do not cluster their chromosomes at the pronuclear interface, and may underlie the lower aneuploidy rates in mouse embryos compared to human and bovine embryos (Destouni et al., 2016; Lee and Kiessling, 2017; Lightfoot et al., 2006). Additionally, chromosome clustering may be important for other processes in early embryo development, such as the establishment of new topologically associating domains (Borsos et al., 2019; Chen et al., 2019) and embryonic genome activation (Li et al., 2013).

Our results suggest that clustering of nucleoli in human zygotes could be used as a proxy for efficient chromosome clustering and improved embryo development, though further studies using a larger dataset of human embryos are required (Fulka et al., 2015; Guerif et al., 2007; James et al., 2006; Swain, 2013). This approach would be of particular interest to *in vitro* fertilization clinics in countries such as Germany that freeze zygotes before the pronuclei break down.

## Acknowledgements

We thank L. Pieper and M.W. Kriesten (Westfleisch, Hamm, Germany) for enabling this research by dissecting and donating bovine ovaries; the MPI-BPC drivers for collecting ovaries from Hamm; M. Daniel for assistance in coordinating ovarian collections; J. Uraji and K. Menelaou (MPI-BPC) for help in providing the human embryo videos; C. So (MPI-BPC) for technical advice; S. Munro (LMB, Cambridge, UK), A. Musacchio (MPI-Molecular Physiology, Dortmund, Germany), J. Ellenberg (EMBL, Heidelberg, Germany), and X.W. Wang (National Cancer Institute/NIH, Bethesda, USA) for plasmids; Z. Holubcová (Department of Histology and Embryology, Masaryk University) for discussions about the division of human zygotes; the live-cell imaging facility and P. Lénárt (MPI-BPC) for discussion and support; C. Thomas (MPI-BPC), L. Wartosch (MPI-BPC), and Life Science Editors for critical comments on the manuscript; members of the Schuh lab for discussions. The research leading to these results has received financial support from Max Planck Society and Deutsche Forschungsgemeinschaft (Multiscale Bioimaging Excellence Cluster and Leibniz Prize to M.S.). The authors declare no competing financial interests.

## Authors contributions

T.C and M.S. conceived the study; T.C. and M.S. designed experiments; T.C. performed all experiments; T.C., A.Z.P., and M. S. designed methods for data analysis; A.Z.P. wrote all in-house–developed scripts and plugins; T.C. and A.Z.P. analyzed the data: P.A., A.L.H., and H.N. supervised the establishment of the experimental system; M.B. and K.E. supervised the work at the IVF clinic; C.B. recorded the human embryos videos; T.C. and M.S. wrote the manuscript and prepared the figures with input from all authors; and M.S. supervised the entire study.

## Materials and methods

### Bovine zygote production

Bovine ovaries were obtained from a local abattoir and transported to the laboratory in a thermo-flask within 3 hours of retrieval. Oocyte isolation, culture, and fertilization were performed using the IVF Bioscience media suite and the manufacturer protocol, with small changes. In brief, cumulus-oocyte complexes (COCs) were aspirated from antral follicles using an 18-gauge needle mounted on a 1 mL disposable syringe. The aspirated follicular fluid was transferred to a 50 ml falcon tube containing 140 µl of 5000 IU/ml Heparin (Merck Millipore). COCs were allowed to sediment and then washed extensively with pre-warmed TCM199 (HEPES-buffered medium 199; Sigma M2520) supplemented with 0.05 g/ml Gentamycin Sulphate (Roth 0233), 1 mM Na-Pyruvate (GIBCO 11360-039), 0.022 g/ml NaHCO3 (S5761-500G), and 5% FBS (GIBCO 16000-044). Only fully-grown oocytes with a homogeneous cytoplasm and at least 3-5 complete layers of compact cumulus cells were selected for the experiments. COCs were washed and transferred to pre-warmed and equilibrated BO-IVM media (IVF Biosciences) and incubated at 38.8°C (5% CO2). After 14 hours, COCs were partially denuded using a transfer pipette with a 175 µm tip (Origio MXL3-175) in warm TCM199 media. 6 hours later (20 hours after COCs retrieval), COCs were washed into pre-warmed and equilibrated BO-IVF media (IVF Biosciences). Insemination was performed by adding 1*106 spermatozoa to a maximum of 40 COCs in a final volume of 500 µl BO-IVF media. Spermatozoa were purified from frozen bull semen obtained from a bull of proven fertility (Bernal-Ulloa et al., 2016). Frozen semen was thawed, resuspended in 3 ml of pre-warmed BO-Semen media (IVF Bioscience) and centrifuged for 5 min at 300 g. The pellet was resuspended in 2 ml of pre-warmed BO-Semen media and ccentrifuged for 5 min at 300 g. The supernatant was discarded leaving 300-500 µl and sperm cells were gently resuspended and counted using a Bürker chamber. Insemination was performed at 38.8°C (5% CO2) for 7 to 18 hours. For short inseminations (7-10 hours), spermatozoa concentration was doubled. Zygotes were retrieved 7 to 18 hours after insemination and gently denuded of cumulus cells and spermatozoa using a transfer pipette with a 135 µm tip (Origio MXL3-135) in pre-warmed TCM199 media. To improve imaging, zygotes were transferred to a 2 ml tube containing 500 µl of warm TCM199 media and centrifugated for 3 min at 9000 g in a pre-warmed rotor. After spinning, zygotes were washed into pre-warmed and equilibrated BO-IVC media (IVF Biosciences) and incubated at 38.8°C (5% CO2 + 6% O2).

### Human embryos

All human embryos used in this study were part of the routine IVF treatment after having obtained fully informed patient consent at Bourn Hall Clinic (Cambridge, UK), under Human Fertilization and Embryology Authority (HFEA) license for Center 0100.

### Microinjection of bovine oocytes and zygotes

Bovine oocytes and zygotes were microinjected with 4 pl of mRNAs as previously described (Schuh and Ellenberg, 2007). Bovine oocytes were injected between 14 and 20 hours after onset of maturation. Bovine zygotes were injected between 8 and 16 hours post insemination. mRNAs were microinjected at the following needle concentrations: mClover3-MAP4-MTBD at 200 ng/µl, miRFP-MAP4-MTBD at 300 ng/µl, H2B-miRFP at 70 ng/µl, H2B-mScarlet at 60 ng/µl, mScarlet-hCenpC at 41 ng/µl, GST at 1964 ng/µl, mClover3-bKASH5-DN at 1950 ng/µl, GST-bKASH5-DN at 1951 ng/µl, bElys-mClover3 at 432 ng/µl, H2B-mClover3 at 120 ng/µl, POM121-mScarlet at 50 ng/µl, and mClover3-hNPM1 at 24 ng/µl.

For protein injections, zygotes were microinjected with 14 pl. BSA and P150-CC1 were injected at a needle concentration of 20 µg/µl, resulting in a protein concentration in zygotes of 4.7 mM and 7.1 mM, respectively. Protein injection buffer was PBS supplemented with 0.03% NP40 and 70 kD Dextran 647 to select for injected cells.

### Expression constructs, messenger RNA (mRNA) synthesis, protein expression and purification

All mRNAs were synthesized and quantified as previously described (So et al., 2019). mClover3-MAP4-MTBD, H2B-miRFP, and H2B-mScarlet mRNAs were synthesized from previously published constructs (So et al., 2019). The following plasmids were generated specifically for this study by subcloning the previously published sequences into a pGEMHE plasmid: GST (Panic et al., 2003), mRFP-MAP4-MTBD (So et al., 2019), H2B-mClover3 (So et al., 2019); mScarlet-hCenpC (Klare et al., 2015), Pom121-mScarlet (Beaudouin et al., 2002), mClover3-hNPM1 (Wang et al., 2005). mClover3-KASH5-DN, GST-KASH5-DN, and mClover3-bElys constructs were cloned from bovine fibroblast or bovine oocyte cDNA libraries made using a SensiFAST cDNA synthesis kit (Bioline, BIO-65053). The primers (KASH5-DN cloning, sense and antisense, Elys cloning sense and antisence) were used to clone the bovine KASH5-DN and bovine Elys into a pENTR/D-TOPO vector (Invitrogen).

His-P150-CC1 (Courtois et al., 2012) was expressed in and purified from BL21(DE3)pLysS competent cells (Promega) as previously described (Courtois et al., 2012) with some modifications. Briefly, recombinant proteins were purified using Ni-NTA Agarose (Qiagen 30210). Buffer was exchanged by dialysis to PBS. Protein was concentrated using an Amicon 10 kD filter column (Merck UFC801008D) and purity tested by SDS-PAGE.

### Drug addition

Nocodazole (Sigma-Aldrich M1404) was diluted freshly in hybridoma grade DMSO (Sigma-Aldrich D2650) to make a 20 mM stock and was added to zygotes to a final concentration of 20 µM, at least 9 hours before nuclear envelope breakdown.

### Immunofluorescence

Zygotes were fixed in 100 mM HEPES (pH 7.0, titrated with KOH), 50 mM EGTA (pH 7.0, titrated with KOH), 10 mM MgSO_4_, 2% methanol-free formaldehyde and 0.5% triton X-100 at 37°C for 25 min. Fixed zygotes were extracted in PBS with 0.5% triton X-100 (PBST) overnight at 4°C and blocked in PBST with 5% BSA (PBST-BSA) for 6 hours at room temperature. Primary antibody incubations were performed in PBST-BSA overnight at 4°C at the concentration listed in the following paragraph. Secondary antibody incubations were performed in PBST-BSA for 1 hour at room temperature at 20 μg/ml.

Primary antibodies used were human anti-centromere antibody (ACA) at 1:250 dilution (Antibodies Incorporated #15-234), rat anti-Nup98 at 1:50 (Abcam #ab50610), mouse anti-NPC/MAb414 at 1:100 (Covance #MMS-120P), rat anti-α-tubulin at 1:1000 (AbD Serotec - #MCA78G), rabbit anti-β-8-tubulin at 1:500 (Sigma-Aldrich #SAB2700070), mouse anti-Trf1 at 1:250 (Alpha diagnostic international #TRF12-S), mouse anti-Histone at 1:100 (Merck #MAB3422), mouse anti-γ-tubulin at 1:250 (Sigma-Aldrich # T5326), rabbit anti-Sun1 at 1:100 (CST # 8886), rabbit anti-Lamin B1 at 1:100 (Abcam #ab16048) and mouse anti-Lamin A/C at 1:50 (Sigma-Aldrich #MABT1340).

Secondary antibodies used were Alexa Fluor 405-, 488-, 568- or 647-conjugated anti-human IgG, anti-mouse IgG, anti-mouse IgM, anti-rabbit IgG, or anti-rat IgG all raised in donkey or goat (Thermo Fisher Scientific). DNA was stained with DAPI at final concentration of 20 µg/ml (Thermo Fisher Scientific).

### Confocal and super-resolution microscopy

For confocal imaging, oocytes were imaged in 20 µl of BO-IVC medium (for live zygotes) or PBS with 1% polyvinylpyrrolidone (PVP) (for fixed oocytes) under paraffin oil in a 35 mm dish with a #1.0 coverslip. Images were acquired with LSM800, LSM880, or LSM900 confocal laser scanning microscopes (Zeiss) equipped with an environmental incubator box and a 40x C-Apochromat 1.2 NA water-immersion objective. A volume of 65 µm × 65 µm × 60 µm centered on the chromosomes was typically recorded. If full oocytes were imaged, then we used a volume of 100 µm × 100 µm × 72.5 µm centered on the zygote center. mClover3 was excited with a 488 nm laser line and detected at 493 - 571 nm. mScarlet was excited with a 561 nm laser line and detected at 571 - 638 nm. miRFP was excited with a 633 nm laser line and detected at 638 - 700 nm. Images of the control and experimental groups were acquired under identical imaging conditions on the same microscope. For some images, noise was reduced with a Gaussian filter in ZEN (Zeiss). Airyscan images were acquired using the Airyscan module on LSM800, LSM880, or LSM900 confocal laser scanning microscopes (Zeiss) and processed in ZEN (Zeiss) after acquisition. Care was taken that the imaging conditions (laser power, pixel-dwell time and detector gain) did not cause phototoxicity (for live imaging), photobleaching or saturation. Selective photobleaching experiments were performed on zygotes expressing H2B-mClover3 and H2B-mScarlet. In experiments where zygotes expressed H2B-mScarlet, bleaching was done once chromosomes were already distributed on the nuclear surface or right after NEBD. Bleaching was performed using the bleaching tool of ZEN (Zeiss) with 60% 561 laser power, scan speed 4, and three repeats. Z-projections were done by maximum intensity projections of the indicated Z-stacks.

### Human embryo imaging

Time-lapse images of human embryos were recorded using a GERI (Genea Biomedx) system as part of routine IVF treatment with patient consent at Bourn Hall Clinic, under Human Fertilization and Embryology Authority (HFEA) license for Center 0100. Imaging was offered to patients for the purpose of selecting embryos with optimal implantation potential. Videos were recorded using the. Data for the present study was obtained via retrospective analysis of archived anonymized records.

### Statistical analysis

Statistical significance based on paired or unpaired, two-tailed Student’s t-test (for absolute values) and two-tailed Fisher’s exact test (for categorical values) were calculated in Prism (GraphPad). All box plots show median (horizontal black line), mean (small black squares), 25^th^ and 75^th^ percentiles (boxes), 5^th^ and 95^th^ percentiles (whiskers) and 1^st^ and 99^th^ percentiles (crosses). All data are from at least three independent experiments. *P* values are indicated.

### Image analysis

DNA-Nuclear Pore Complex co-localization analysis was performed using Imaris version 9.2.1 (Bitplane). We used the Imaris spot tool to create a sphere for each nucleus. The subsequent analysis was performed within the spheres. Using the surface tool on the NPC channel, we identified the membrane regions containing nuclear pores. With the Imaris co-localization analysis tool we computed the Pearson correlation coefficient between the DNA and NPC signals.

The quantification of microtubule length was performed using Imaris spot tool. For microtubules originating from the centrosome we marked the centrosome and the opposite microtubule end and measured the distance between the two points. The microtubule length at metaphase was measured from astral microtubules. For this quantification we used zygotes fixed at metaphase and stained for microtubules (α-tubulin) and DNA (DAPI).

The quantification of bleached and not bleached lagging chromosomes originated in zygotes having pronuclei with equally condensed chromosomes (Fig. S4E) was performed using Imaris. The Imaris surface detection tool was applied on the unbleached channel (laser 488 – H2B-mClover3) to identify bulk chromatin and individual chromosomes. For each zygote, we computed the average ratio of the H2B-mClover3 and the H2B-mScarlet signals for three surfaces with bleached DNA (*B*) and for three surfaces with unbleached DNA (*U*). We also computed the average ratio of the H2B-mClover3 and the H2B-mScarlet for each lagging chromosome (*LC*). The quantities *B* and *U* were calculated just after the bleaching. Lagging chromosomes with |*B* - *LC*| < |*U* - *LC*| were scored as bleached (clustered). Lagging chromosomes with |*B* - *LC*| > |*U* - *LC*| were scored as unbleached (distal).

The quantification of bleached and not bleached lagging chromosomes originated in zygotes having a pronucleus with uncondensed chromosomes (Fig. S4, B and C) was performed using Imaris. The Imaris surface detection tool was applied on the unbleached channel (laser 488 – H2B-mClover3) to identify bulk chromatin and individual chromosomes. For each zygote, we computed the ratio of the H2B-mClover3 and the H2B-mScarlet signals in the bleached pronucleus (*B*) and in the unbleached pronucleus (*U*). We also computed the average ratio of the H2B-mClover3 and the H2B-mScarlet for each lagging chromosome (*LC*). The quantities *B* and *U* were calculated just after the bleaching. Lagging chromosomes with |*B* - *LC*| < |*U* - *LC*| were scored as bleached (clustered). Lagging chromosomes with |*B* - *LC*| > |*U* - *LC*| were scored as unbleached (uncondensed).

The DNA distribution indexes (nuclear and surface occupancy indexes and the inner, outer and maximal polarity indexes) were computed using Fiji (Rueden et al., 2017; Schindelin et al., 2012)(IJ2 v2.0.0-rc-71, IJ1 v1.52r) and MATLAB (v2018a, The MathWorks, Inc.). A sphere is used to approximate each pronucleus, the center coordinates ***S*** *=* (*S*_*X*_, *S*_*Y*_, *S*_*Z*_) and radius *R* were calculated from points placed manually in the image (*Fit Sphere*, BoneJ2 v6.1.1) (Doube et al., 2010). We developed the ImageJ plugin *liveim-drawspheres* to compute pronuclei masks using ***S*** and *R* and a maximal polarity direction per nucleus. For this, Gaussian filtered images of the DNA signal using [*σ*_*X*_, *σ*_*Y*_, *σ*_*Z*_] equal to [8, 8, 0], [16, 16, 0], [16, 16, 1], [32, 32, 0], [32, 32, 1], and [32, 32, 2] pixels are calculated. The DNA intensity image and the pronucleus mask is used to compute the coordinates, ***CM***, of the center of mass for the intensity distribution and for extended maxima regions (MorphoLibJ v1.4.1) (Legland et al., 2016). For each center of mass, the mean intensity in three equally spaced sphere sections along the vector ***p*** = (***CM*** − ***S***) is computed. This operation is performed for the raw and Gaussian filtered DNA images and the orientation with the highest normalized mean fluorescence intensity in the first region (the region facing the center of mass) approximates the direction of maximal polarity ***p***_***max***_. The ImageJ plugin also computes distance transform for each pronucleus and defines sphere sections oriented along the line connecting the center of the nuclei. These are used to compute the inner and outer polarity indexes.

In the last part of the workflow, a MATLAB script is used to extract the relevant quantities per pronucleus. We first smoothed the DNA signal using a 2D median filter (3×3 pixels) and a Gaussian filter (*σ* = 1 pixel). To minimize the contribution of unbound H2B-fluorescent protein, we derived a mask for the DNA, *D*, using an Otsu’s threshold *thr(pre-NEBD)* (*multithresh*). The pixel values outside *D* are set to 0. The ratio of number of pixels in *D* to the whole nucleus gives the:

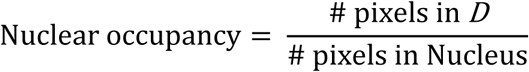

The surface occupancy is calculated from:

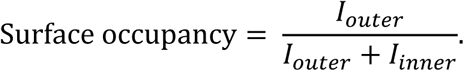

Where *I*_*outer*_ and *I*_*inner*_ are, respectively, the mean intensities in the region adjacent to the nuclei surface and inside the nucleus. The distance transform is used to define the region boundaries so to have equal number of pixels in both regions. For a perfect sphere this is ∼ 0.8 *R*. The sphere sections have been chosen to have equal heights *h* = 2*R*/3, yielding equal outer surface areas, i.e. the area without the base of the sections. For sphere sections oriented toward ***p***_***max***_ we computed the mean fluorescence intensity *I*_*k*_ and normalized it to obtain

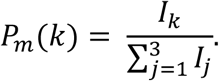

Similarly, we computed mean intensities for sections oriented toward the opposite nucleus to obtain *P*_*o*_(*k*), *k* =1, 2, 3. Where section 1 is the section in the direction of maximal polarity/toward the other nucleus, 2 the central section, and 3 the section which is oriented in the opposite direction. The maximal polarity is given by *P*_*m*_(1) the inner and outer polarities are given by *P*_*o*_(1) and *P*_*o*_(3), respectively.

For time lapse data (Fig. 1C) the threshold *thr*(t) to segment the DNA is adapted per time point. We computed the total intensity in *D* and adapted the threshold so that the total intensity remains within 10% of the pre-NEBD value. Variation in intensity during migration, is accounted for by computing a correction factor from the total non-segmented DNA intensity at pre-NEBD and the respective time point.

Scripts and plugins are available at link: https://gitlab.gwdg.de/schuh-meiosis (*active after manuscript acceptance for publication, currently attached as additional material*).

## Supplementary figure legends

**Figure S1.**
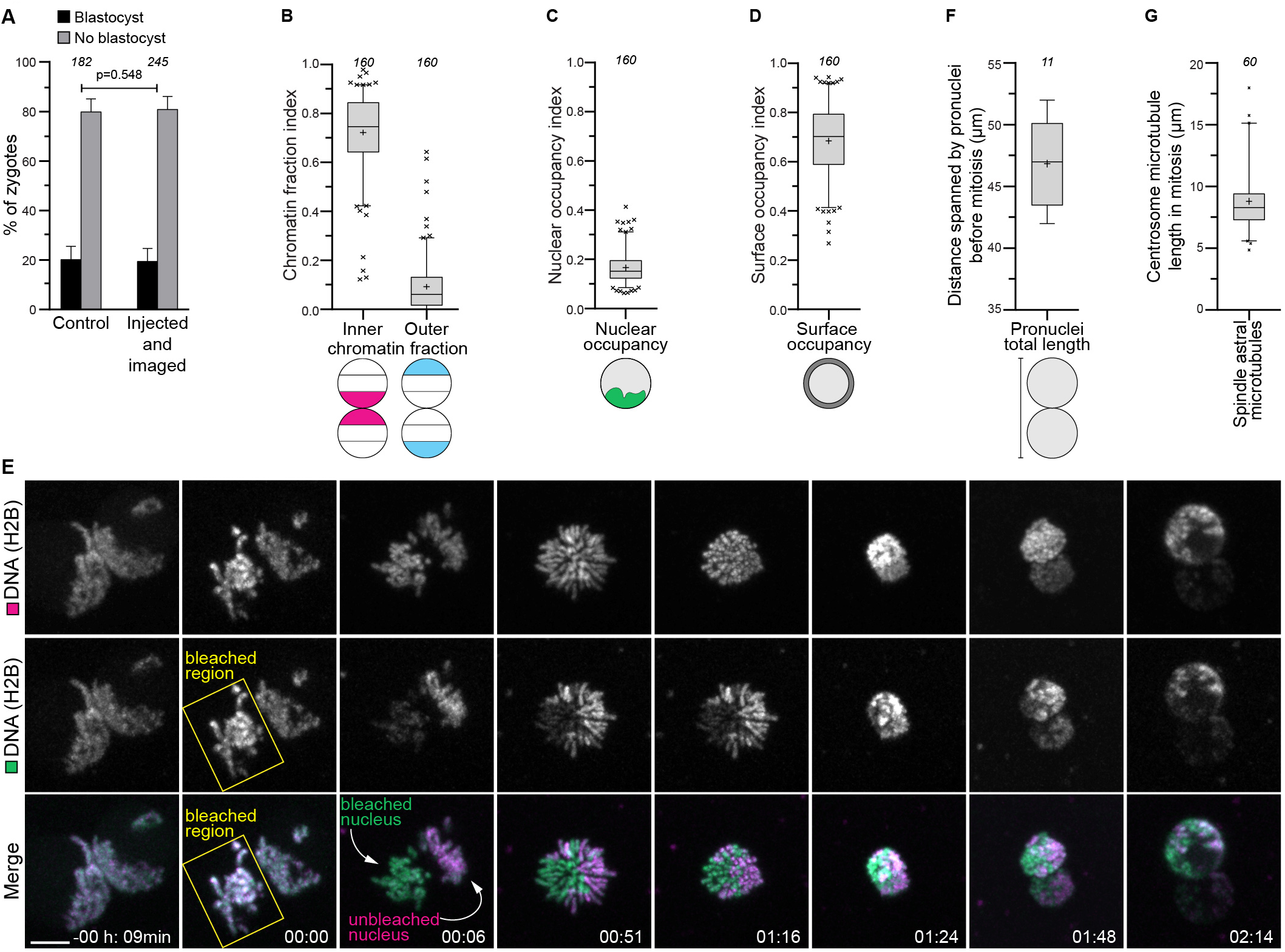
Maternal and paternal chromosomes cluster at the pronuclear interface, related to Figure 1. (A) Cow zygotes non-injected and non-imaged or injected and imaged that developed into blastocysts or developed abnormally. (B) Inner and outer chromatin fraction indices before nuclear envelope breakdown. (C) Nuclear occupancy index before nuclear envelope breakdown. (D) Surface occupancy index before nuclear envelope breakdown. (E) Representative stills from time-lapse movie of a zygote where the H2B-mScarlet chromatin signal in the left pronucleus had been bleached upon nuclear envelope breakdown. Green, DNA (H2B-mClover3). Magenta, DNA (H2B-mScarlet). Bleached region is indicated by a yellow rectangle. The bleached DNA has a green signal, while the unbleached DNA is visible both in green and magenta. Time, hours:minutes, 00:00 is nuclear envelope breakdown. Z-projections, 7 sections every 1.76 µm. (F) Distance spanned by pronuclei before nuclear envelope breakdown. (G) Centrosome microtubule length at metaphase. Data are from five (A), eleven (B, C, D), and two (F) independent experiments. Data in (G) are from three zygotes obtained from two independent experiment. The number of analyzed zygotes (A, F), pronuclei (B, C, D), or microtubules (G) is specified in italics. Indicated *p*-value calculated using Fisher’s exact test. Scale bars, 10 µm.

**Figure S2.**
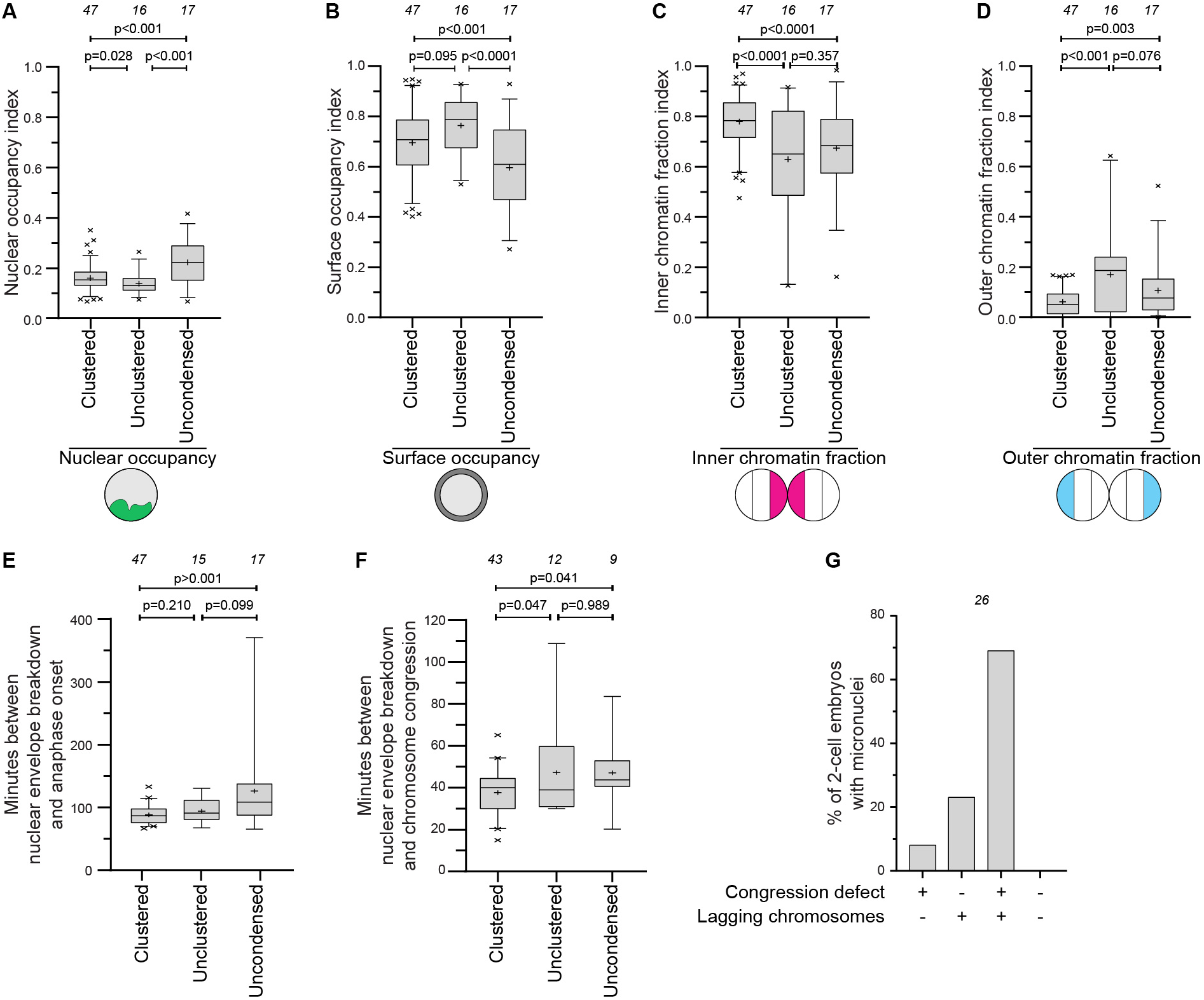
Defective clustering leads to a delay in chromosome congression, related to Figure 2. (A-D) Quantification of chromatin distribution within pronuclei using the nuclear occupancy index (A), surface occupancy index (B), inner chromatin fraction index (C), and outer chromatin fraction index (D) in indicated groups. (E) Time between nuclear envelope breakdown and anaphase onset in indicated zygote groups. (F) Time between nuclear envelope breakdown and the completion of chromosome congression on the metaphase plate in indicated zygote groups. Zygotes that failed to align all chromosomes before anaphase onset were excluded. (G) 2-cell embryos with micronuclei that displayed chromosome congression defects and/or lagging chromosomes during zygote mitosis. Data are from eleven independent experiments. The number of analyzed pronuclei (A, B, C, D, G) and zygotes (E, F) is specified in italics. Indicated *p*-value calculated using unpaired two-tailed Student’s t test. Scale bars, 10 µm.

**Figure S3.**
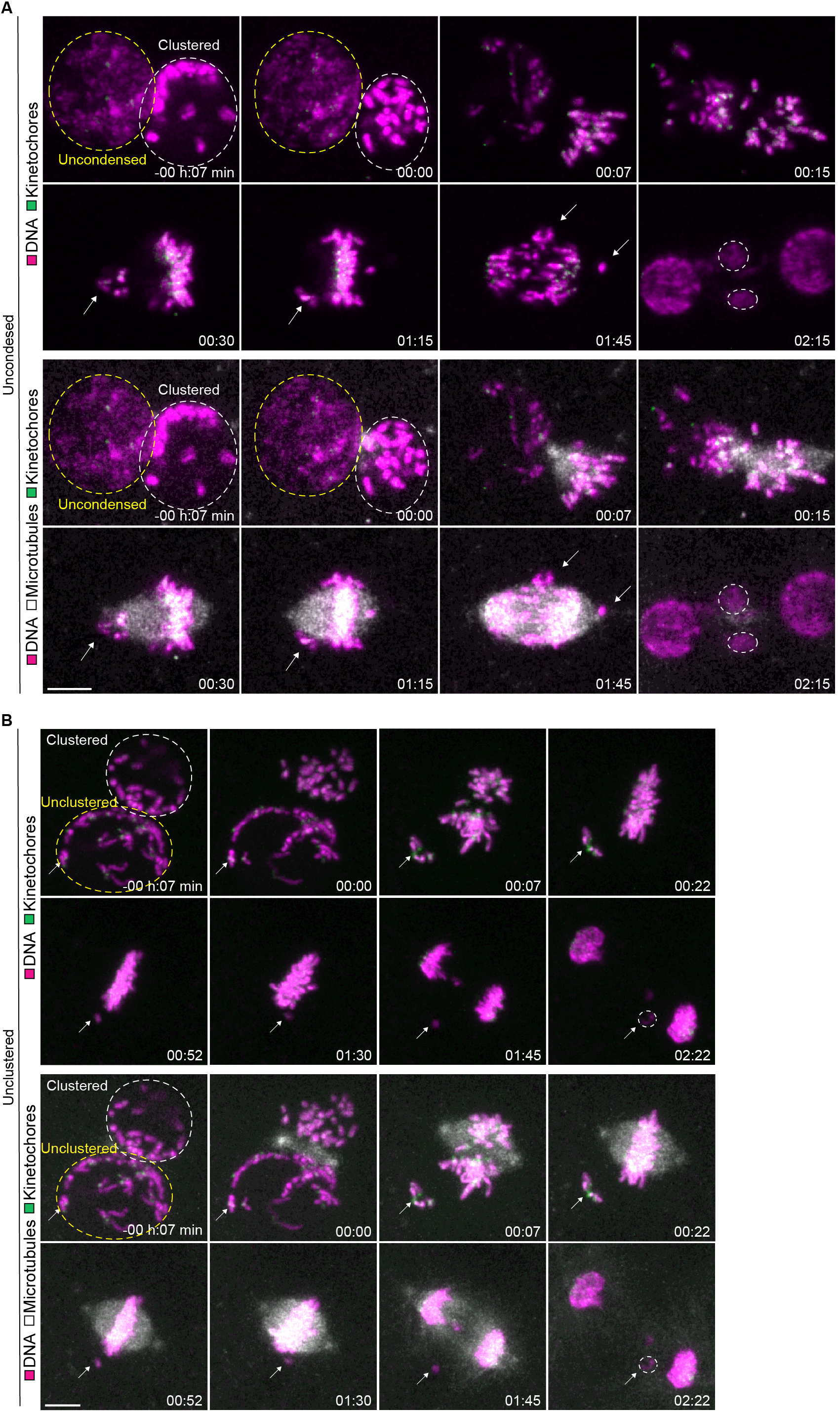
Live kinetochore visualization in zygotes having unclustered and uncondensed chromosomes, related to Figure 2. (A-B) Representative stills from time-lapse movies of zygotes classified as uncondensed (A) and unclustered (B). Gray, microtubules (mClover3-MAP4-MTBD). Magenta, DNA (H2B-miRFP). Green, kinetochores (mScarlet-hCenpC). Dashed lines indicate pronuclei with uncondensed or unclustered chromatin (yellow) and clustered chromatin (white). Arrows point to uncondensed or distal chromosomes that join later the metaphase plate. Several of these chromosomes subsequently form micronuclei, highlighted by white dashed lines. Time, hours:minutes, 00:00 is nuclear envelope breakdown. Z-projections, respectively 4 and 7 sections every 3.08 µm. Scale bars, 10 µm.

**Figure S4.**
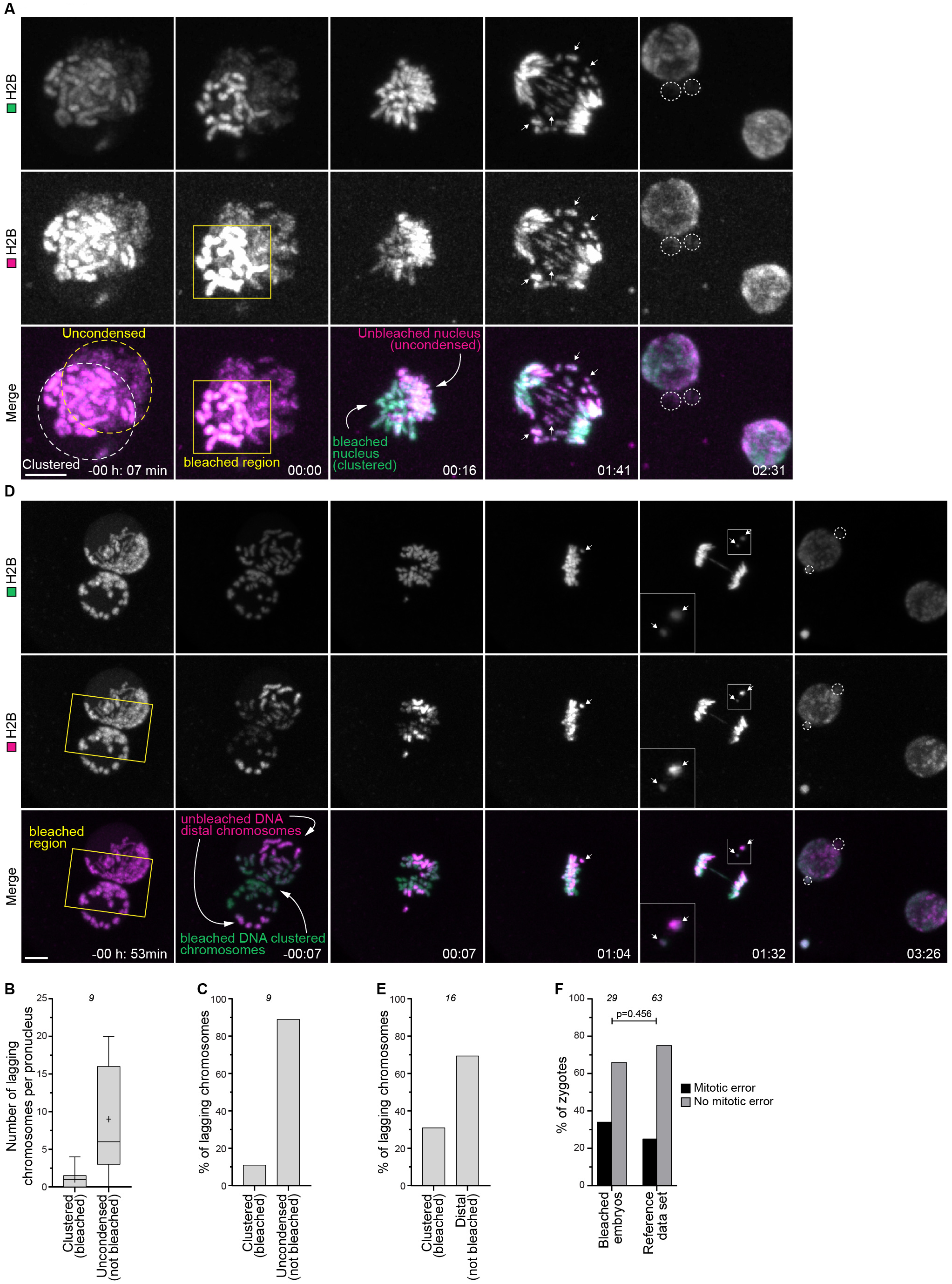
Unclustered chromosomes are more likely to missegregate, related to Figure 2. (A) Representative stills from time-lapse movies of a zygote with one pronucleus having uncondensed chromatin (yellow dashed line). The H2B-mScarlet chromatin signal in the pronucleus with clustered chromatin (left) was bleached upon nuclear envelope breakdown in the region indicated by the yellow rectangle. The bleached DNA has a green signal, while the unbleached DNA is visible both in green and magenta. Lagging chromosomes are magenta and green (arrows), indicating that they originated from the uncondensed pronucleus. Green, DNA (H2B-mClover3). Magenta, DNA (H2B-mScarlet). Dashed lines indicate micronuclei. Time, hours:minutes, 00:00 is nuclear envelope breakdown. Z-projections, 9 sections every 1.76 µm. (B) Number of lagging chromosomes originated from pronuclei with clustered DNA (bleached) or uncondensed DNA (not bleached). (C) Percentage of total lagging chromosomes originated from pronuclei with clustered DNA (bleached) or uncondensed DNA (not bleached). (D) Representative stills from time-lapse movies of a zygote with unclustered chromosomes. The H2B-mScarlet chromatin signal between pronuclei was bleached before nuclear envelope breakdown in the region indicated by the yellow rectangle. Lagging chromosomes are magenta and green (arrows), indicating that they were not bleached and were peripheral chromosomes. Green, DNA (H2B-mClover3). Magenta, DNA (H2B-mScarlet). Dashed lines indicate micronuclei. Time, hours:minutes, 00:00 is nuclear envelope breakdown. Z-projections, 18 sections every 1.76 µm. (E) Percentage of total lagging chromosomes originated from clustered DNA (bleached) or distal DNA (not bleached). (F) Zygotes displaying mitotic errors after bleaching and in reference data set (Fig. 2J, merge of clustered and unclustered groups). Data are from four (B, C, E, F-bleached embryos) and eleven (F-reference data set) independent experiments. The number of analyzed zygotes is specified in italics. Indicated *p*-value calculated using Fisher’s exact test. Scale bars, 10 µm.

**Figure S5.**
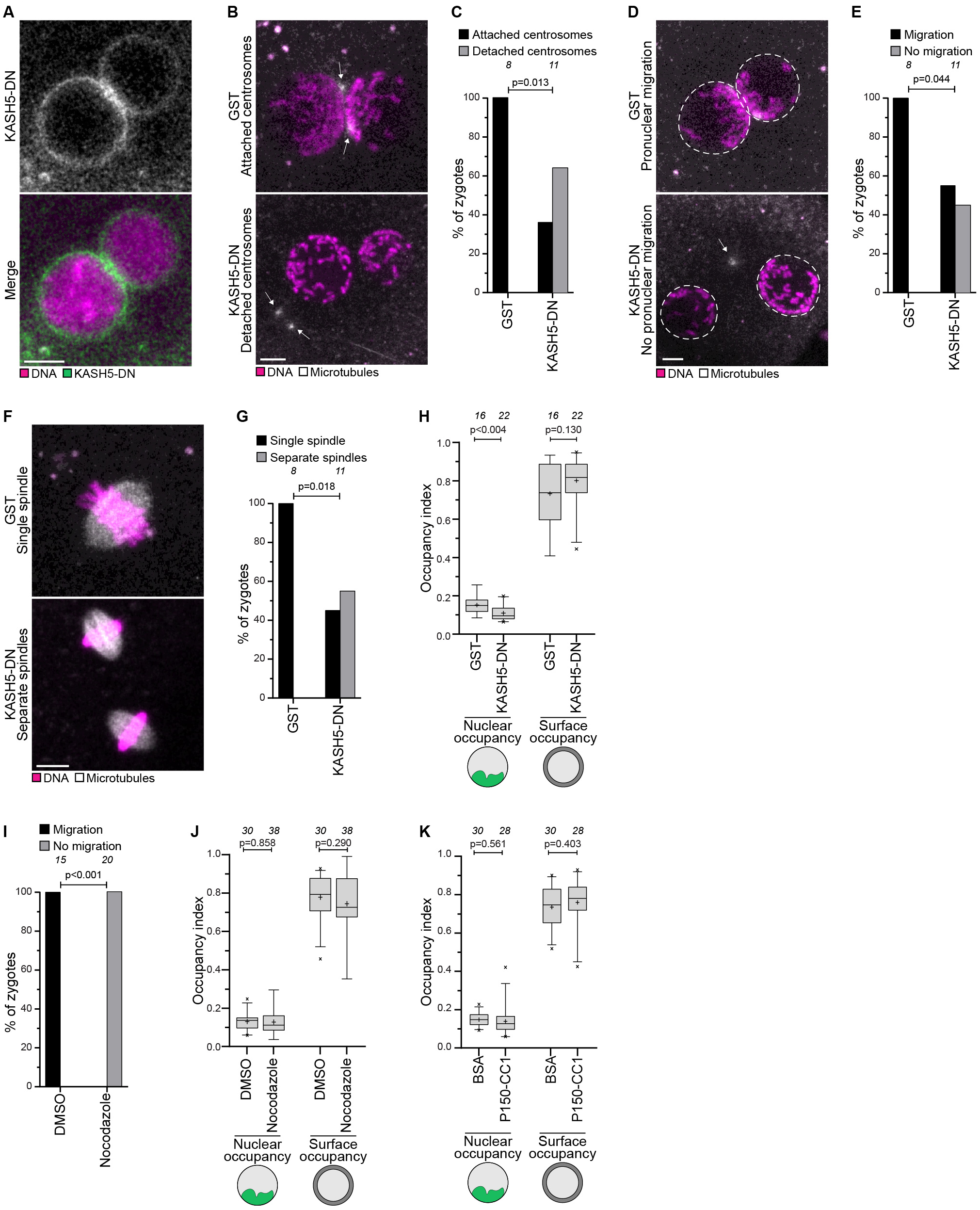
Effects of KASH5-DN, nocodazole, and P150-CC1 treatments, related to Figure 4. (A) Representative image of a zygote expressing mClover3-KASH5-DN (green) and H2B-mScarlet (DNA, magenta). (B-G) Representative images and quantification of zygotes in indicated groups displaying, upon KASH5-DN treatment, detached centrosomes (B and C), pronuclear migration defects (D and E) or separate spindles at anaphase onset (F and G). Gray, microtubules (mClover3-MAP4-MTBD). Magenta, DNA (H2B-mScarlet). Arrows indicate detached centrosomes and dashed lines indicate pronuclear envelopes. Z-projection, respectively 4, 7, 5, 8, 4, and 10 sections every 2.50 µm. (H) Nuclear and surface occupancy indices in zygotes expressing GST or KASH5-DN. (I) Zygotes treated with DMSO or nocodazole that displayed or not pronuclear migration defects. (J) Nuclear and surface occupancy indices in zygotes treated with DMSO or nocodazole. (K) Nuclear and surface occupancy indices in zygotes injected with BSA or P150-CC1. Data are from four (C, E, G, H, K) and six (I, J) independent experiments. The number of analyzed zygotes (C, E, G, I) and pronuclei (H, K, L) is specified in italics. Indicated *p*-values calculated using Fisher’s exact test (C, E, G, I) and unpaired two-tailed Student’s t-test (H, J, K). Scale bars, 10 µm.

**Figure S6.**
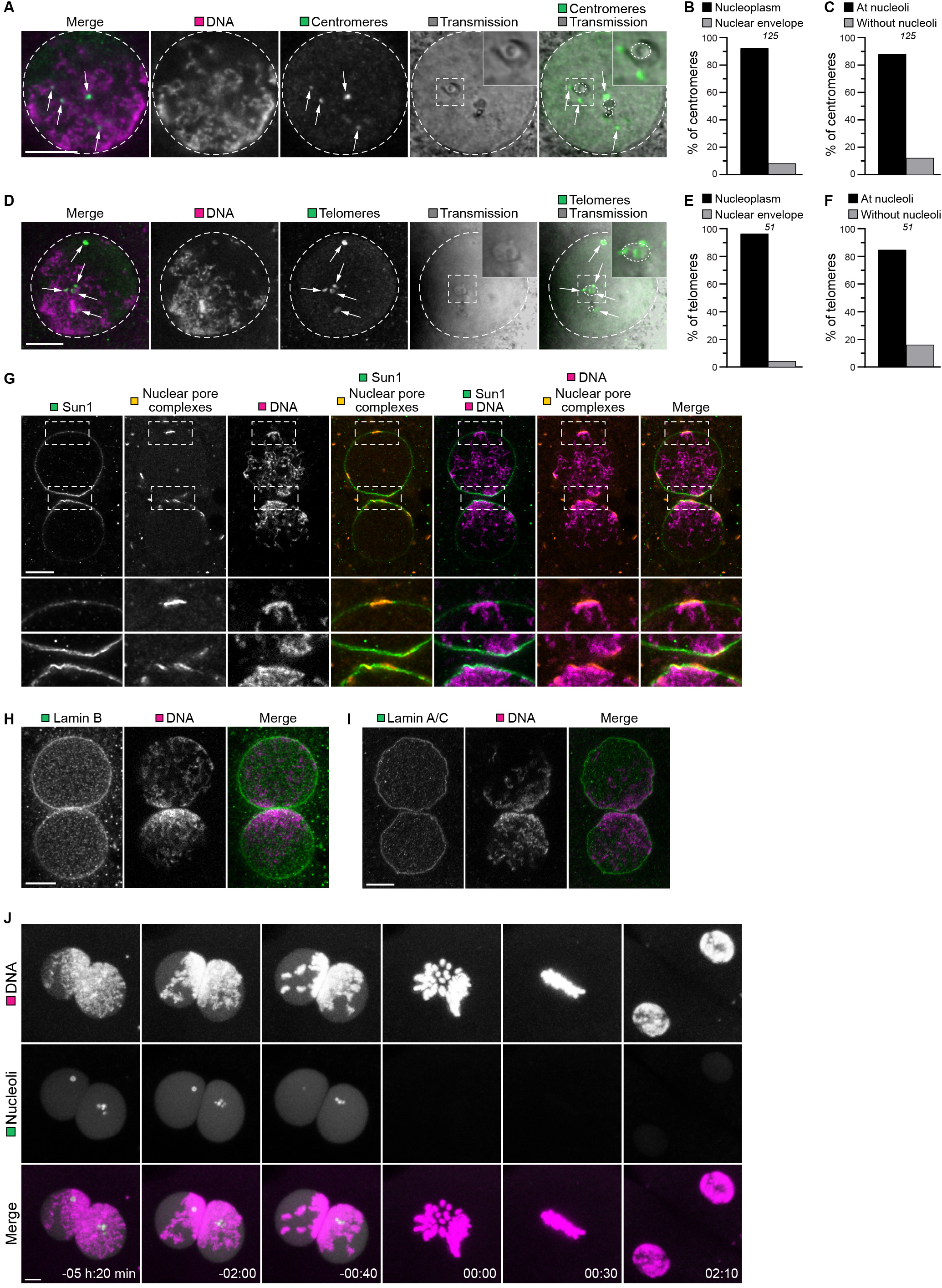
Centromeres, telomeres, Sun1, Lamin B, and Lamin A/C do not cluster at the pronuclear interface. Related to Figures 4-6. (A) Representative immunofluorescence images of centromeres distribution within a pronucleus. Gray, transmission. Magenta, DNA (DAPI). Green, centromeres (ACA). Single section confocal microscopy. Dashed lines indicate pronucleus and nucleoli. Arrows indicate centromeres. Outlined regions magnified in the top right corner. (B) Centromeres in the nucleoplasm or at the nuclear envelope. (C) Centromeres at nucleoli or away from nucleoli. (D) Representative immunofluorescence images of telomere distribution within a pronucleus. Gray, transmission. Magenta, DNA (DAPI). Green, telomeres (Trf1). Single section confocal microscopy. Dashed lines indicate pronucleus and nucleoli. Arrows indicate telomeres. Outlined regions magnified in the top right corner. (E) Telomeres in the nucleoplasm or at the nuclear envelope. (F) Telomeres at nucleoli or away from nucleoli. (G) Representative immunofluorescence images of zygotes stained with Sun1 (Green), Nuclear pore complex (Orange) and DNA (DAPI, Magenta). Outlined regions magnified on the bottom. Single section Airyscan microscopy. (H-I) Representative immunofluorescence images of zygotes stained with Lamin B (H) or Lamin A/C (I) (Green) and DNA (DAPI, Magenta). Respectively, single section Airyscan microscopy and single section confocal microscopy. (J) Representative stills from time-lapse movies of zygotes expressing mClover3-NPM1 (nucleoli, green) and H2B-mScarlet (DNA, magenta). Time, hours:minutes, 00:00 is nuclear envelope breakdown. Z-projections, 8 sections every 2.50 µm. Data are from five embryos (two pronuclei each) generated in a single experiment. The number of analyzed telomeres (B, C) and centromeres (E, F) is specified in italics. Scale bars, 10 µm.

## Supplementary movie legends

**Movie S1. Bovine embryo development from zygote to 8-cell stage, related to Figure 1**.

Time-lapse movies of bovine zygotes undergoing the first cell divisions. Time, hours:minutes, 00:00 is nuclear envelope breakdown.

Part I: A system for high resolution live cell microscopy of early bovine embryo development. Zygote expressing mClover3-MAP4-MTBD (gray, microtubules) and H2B-mScarlet (magenta, DNA). Z-projections, 8 sections every 2.50 µm.

Part II: Chromosome segregation errors during the first mitotic division (mild). Zygote expressing mClover3-MAP4-MTBD (gray, microtubules) and H2B-mScarlet (magenta, DNA). Z-projections, 19 sections every 2.50 µm.

Part III: Chromosome segregation errors during the first mitotic division (severe). Zygote expressing mClover3-MAP4-MTBD (gray, microtubules) and H2B-mScarlet (magenta, DNA). Z-projections, 17 sections every 2.50 µm.

**Movie S2. The maternal and paternal chromosomes occupy partially distinct territories in the metaphase plate of the zygote spindle, related to Figure 1**.

Time-lapse movie of a zygote expressing mClover3-H2B (green, DNA) and H2B-mScarlet (magenta, DNA). Rectangle indicates region bleached using the 561 laser. Time, hours:minutes, 00:00 is nuclear envelope breakdown. Z-projections, 18 sections every 1.76 µm.

**Movie S3. Chromosomes in the pronucleus with uncondensed chromatin lag behind during anaphase, related to Figure 2**.

Time-lapse movies of zygotes having one uncondensed pronucleus undergoing mitosis. Arrowheads indicate chromosomes that fail to align at the metaphase plate, lag after anaphase, and eventually form micronuclei, dotted circles. Time, hours:minutes, 00:00 is nuclear envelope breakdown.

Part I: Zygotes with an uncondensed pronucleus have chromosome segregation errors. Zygote expressing mClover3-MAP4-MTBD (gray, microtubules) and H2B-mScarlet (magenta, DNA). Z-projections, 12 sections every 2.50 µm.

Part II: Kinetochores in the uncondensed pronucleus are less accessible than those at the surface of condensed pronuclei. Zygote expressing mScarlet-hCenpC (green, kinetochores), mClover3-MAP4-MTBD (gray, microtubules) and H2B-miRFP (magenta, DNA). Z-projections, 4 sections every 3.08 µm.

Part III: Chromosome segregation errors originate from uncondensed pronucleus. Zygote expressing mClover3-H2B (green, DNA) and H2B-mScarlet (magenta, DNA). Rectangle indicates region bleached on the condensed pronucleus, using the 561 laser. Z-projections, 10 sections every 1.76 µm.

**Movie S4. Unclustered chromosomes lag behind during anaphase, related to Figure 2**.

Time-lapse movies of zygotes having one unclustered pronucleus undergoing mitosis. Arrowheads indicate chromosomes that fail to align at the metaphase plate, lag after anaphase, and eventually form micronuclei, dotted circles. Time, hours:minutes, 00:00 is nuclear envelope breakdown.

Part I: Zygotes with an unclustered pronucleus and chromosome segregation errors. Zygote expressing mClover3-MAP4-MTBD (gray, microtubules) and H2B-mScarlet (magenta, DNA). Z-projections, 12 sections every 2.50 µm.

Part II: Chromosomes that do not cluster at the pronuclear interface congress late and segregate incorrectly. Zygote expressing mScarlet-hCenpC (green, kinetochores), mClover3-MAP4-MTBD (gray, microtubules) and H2B-miRFP (magenta, DNA). Z-projections, 4 sections every 3.08 µm.

Part III: Chromosomes distal from the pronuclear interface are missegregated. Zygote expressing mClover3-H2B (green, DNA) and H2B-mScarlet (magenta, DNA). Rectangle indicates region bleached at pronuclear interface, using the 561 laser. Z-projections, 17 sections every 1.76 µm

**Movie S5. Centrosomes, microtubules, and dynein pre-unite chromosomes within intact pronuclei, related to Figure 4**.

Time-lapse movies of zygotes treated with different perturbation to affect the function of, respectively, centrosomes, microtubules, and dynein. Time, hours:minutes, 00:00 is nuclear envelope breakdown.

Part I: Centrosomes are required for chromosome clustering. Three zygotes expressing mClover3-MAP4-MTBD (gray, microtubules), H2B-mScarlet (magenta, DNA), and GST (left) or KASH5-DN (middle and right). Arrowheads indicate detached centrosomes before and after nuclear envelope breakdown. Z-projections, 9 sections every 3.08 µm.

Part II: Microtubules are required for chromosome clustering. Two zygotes expressing mClover3-MAP4-MTBD (gray, microtubules) and H2B-mScarlet (magenta, DNA) treated with DMSO (left) or nocodazole (right). Z-projections, respectively 7 and 8 sections every 2.50 µm.

Part III: Dynein is required for chromosome clustering. Two zygotes expressing mClover3-MAP4-MTBD (gray, microtubules) and H2B-mScarlet (magenta, DNA) injected with BSA (left) or P150-CC1 (right). Z-projections, respectively 8 and 13 sections every 2.50 µm

**Movie S6. Annulate lamellae move toward centrosomes at the pronuclear interface, related to Figure 5**.

Time-lapse movie of a zygote expressing mClover3-MAP4-MTBD (orange, microtubules) and POM121-mScarlet (green, nuclear pore complexes). Arrowheads indicate annulate lamellae moving toward the centrosomes. Time, hours:minutes, 00:00 is nuclear envelope breakdown. Single sections confocal microscopy.

**Movie S7. Nuclear pore complexes cluster with chromatin at the pronuclear interface, related to Figure 5**.

Airyscan sections of a zygote stained for microtubules nuclear pore complexes (green, NPC-Mab414), (orange, α-tubulin), and DNA (magenta, DAPI). Outlined regions magnified on the bottom. Centrosomes, nuclear pore complexes, and annulate lamellae are indicated. One section corresponds to 0.18 µm, as indicated.

**Movie S8. Nucleoli as a read out of chromosome clustering and blastocyst development in human embryos, related to Figure 6**.

Time-lapse movies of two human embryos. Left embryo at the zygote stage has clustered nucleoli before nuclear envelope breakdown and it develops into blastocyst. Right embryo has unclustered nucleoli and does not develop into blastocyst.

Time, hours:minutes, 00:00 is nuclear envelope breakdown in the zygote.

